# Sex-specific regulation of the cardiac transcriptome by the protein phosphatase 2A regulatory subunit B55α

**DOI:** 10.1101/2024.05.05.592558

**Authors:** Nicola M Sergienko, Adam J Trewin, Helen Kiriazis, Antonia JA Raaijmakers, Daniel G Donner, Victoria C Garside, Kelly A Smith, James R Bell, Kimberley M Mellor, Lea MD Delbridge, Julie R McMullen, Kate L Weeks

**Affiliations:** Baker Heart and Diabetes Institute, Melbourne, Victoria, Australia; Department of Diabetes, Central Clinical School, Monash University, Clayton, Victoria, Australia; Department of Anatomy and Physiology, The University of Melbourne, Melbourne, Victoria, Australia; Department of Cardiometabolic Health, The University of Melbourne, Melbourne, Victoria, Australia; Department of Microbiology, Anatomy, Physiology & Pharmacology, La Trobe University, Melbourne, Victoria, Australia; Department of Physiology, University of Auckland, New Zealand; Department of Cardiovascular Research, Translation and Implementation, La Trobe University, Melbourne, Victoria, Australia; Monash Alfred Baker Centre for Cardiovascular Research, Monash University, Melbourne, Victoria, Australia

**Keywords:** PP2A, sex differences, development, ageing, ECM

## Abstract

Protein phosphatase 2A (PP2A) regulatory subunit B55α has been implicated in the transcriptional regulation of cardiac growth and fibrosis by suppressing HDAC5/MEF2 signalling in cardiomyocytes. We created and characterised two mouse models with global or cardiomyocyte-specific disruption of the gene encoding B55α (*Ppp2r2a*) to conduct the first detailed exploration of B55α in the heart. Global homozygous B55α knockout mice died *in utero*, while heterozygous mice had thinner left ventricular walls at 12 months, an effect more pronounced in males. At 10-12 weeks of age, cardiomyocyte-specific B55α knockout mice displayed normal cardiac morphology with increased left ventricular collagen deposition, identifying B55α as a negative regulator of cardiac fibrosis. Gene expression analyses revealed extensive remodelling of the cardiac transcriptome in male but not female mice, identifying a sexually dimorphic role for B55α in cardiac transcriptional regulation. These findings provide a basis for future work investigating B55α in cardiac stress settings.

## 2. Introduction

Protein phosphatase 2A (PP2A) comprises a family of serine/threonine phosphatases with important roles in cardiac physiology and pathophysiology.^1^ PP2A exists in two forms within the cell: as a heterodimer (the ‘core enzyme’) or as heterotrimers (active holoenzymes). The core enzyme consists of a catalytic ‘C’ subunit and a scaffolding ‘A’ subunit, each encoded by two isoforms (Cα/β and Aα/β). Association of the core enzyme with regulatory ‘B’ subunit isoforms gives rise to distinct PP2A holoenzymes, with different subcellular localisation patterns, substrate specificities and physiological functions.^2^ More than 20 B subunit isoforms and splice variants have been identified, which can be grouped into four subfamilies: B/B55/PR55, B’/B56/PR61, B”/PR72 and B”’/striatins. Detailed study of PP2A B subunits in the heart is largely limited to a single isoform: B56α of the B’/B56 subfamily. *In vitro* and *in vivo* investigations have revealed information about its localisation in cardiomyocytes, key adaptor protein interactions and its role in cardiac contractility.^3–8^ Investigation of other B subunit isoforms is critical for unravelling the complexity of PP2A signalling in the heart.

B55α of the B/B55 subfamily is a ubiquitously expressed PP2A regulatory subunit, with a high level of conservation between species at both the gene and protein level.^9–11^ Its function in the heart has not been specifically investigated. We previously showed that B55α targets components of the PP2A core enzyme to histone deacetylase 5 (HDAC5) in adult rat ventricular myocytes.^12^ This association increased with acute β-adrenergic receptor stimulation, leading to HDAC5 dephosphorylation, nuclear accumulation and repression of myocyte enhancer factor-2 (MEF2) transcriptional activity.^12^ MEF2 transcription factors are essential for cardiogenesis and are critical regulators of cardiac hypertrophy and fibrosis in settings of cardiac stress/injury.^13–16^ Studies in non-cardiac cell lines and tissues have reported that B55α dephosphorylates several other proteins with known roles in cardiac development and hypertrophy. These include Akt and FoxO1,^17,18^ key regulators of exercise-induced physiological cardiac hypertrophy,^19–21^, and β-catenin,^22^ a transcription factor involved in second heart field formation during embryogenesis and pathological cardiac hypertrophy induced by angiotensin II infusion or pressure overload.^23–25^ Therefore, we hypothesised that B55α regulates cardiac development and hypertrophy *in vivo*.

As a first step to investigate B55α in the heart, we generated mice with global or cardiomyocyte-specific deletion of the gene encoding B55α (*Ppp2r2a*) and characterised the impact of reduced B55α expression on cardiac morphology, function and gene expression under basal conditions (i.e. in the absence of a physiological/pathological stress stimulus). Our findings validate previous work showing that B55α is critical for embryogenesis^11,26^ and provide the first characterisation of a B/B55 family member in the adult mouse heart. We demonstrate that cardiomyocyte B55α is not required for cardiac growth during embryonic development or the postnatal period. Reduced B55α expression did, however, lead to thinning of the left ventricular walls with ageing in male mice heterozygous for B55α (global knockout model). We also report a sex-specific role for B55α, with loss of cardiomyocyte B55α having a significant impact on the cardiac transcriptome in male mice, but not in female mice. These studies lay the foundation for future work investigating the role of B55α in cardiometabolic disease settings.

## 3. Methods

### 3.1. Experimental animals

<u>Model 1</u>: C57BL/6N mice heterozygous (Het) for a *Ppp2r2a^tm2a(KOMP)Wtsi^* ‘knockout-first’ allele (RRID: MMRRC:060623-UCD; see **Fig 1a**) were generated by the Australian Phenomics Network (Monash University, Melbourne, Australia). <u>Model 2</u>: To generate cardiomyocyte-specific *Ppp2r2a* knockout (cKO) mice, mice heterozygous for the knockout-first allele were crossed with hemizygous transgenic mice expressing Flp recombinase (Flp^Tg/0^ mice; RRID: IMSR_JAX:003800; see **Fig 1a**). This removed the *lacZ* reporter and neomycin selection cassette and converted *Ppp2r2a^tm2a(KOMP)Wtsi^* to a floxed *Ppp2r2a* allele. Next, offspring hemizygous for Flp and heterozygous for the floxed *Ppp2r2a* allele (Flp^Tg/0^ Ppp2r2a^L/+^) were crossed to remove the Flp transgene. Flp-negative floxed *Ppp2r2a* mice (Flp^0/0^ Ppp2r2a^L/L^) were then mated with hemizygous transgenic mice expressing Cre recombinase under the control of the αMHC promotor (αMHC-Cre^Tg/0^ mice, FVB background).^20,27^ Importantly, Cre expression in this strain does not lead to cardiotoxicity.^20,28^ Cre^Tg/0^ Ppp2r2a^L/+^ and Cre^0/0^ Ppp2r2a^L/+^ mice were mated to produce the following 6 cardiomyocyte-specific genotypes: Cre^Tg/0^ Ppp2r2a^L/L^ (B55α knockout, cKO), Cre^0/0^ Ppp2r2a^L/L^ (floxed control, FC), Cre^Tg/0^ Ppp2r2a^L/+^ (B55α heterozygote, cHET), Cre^0/0^ Ppp2r2a^L/+^ (half-floxed control), Cre^Tg/0^ Ppp2r2a^+/+^ (Cre control), Cre^0/0^ Ppp2r2a^+/+^ (wildtype control). FC and cKO mice for the RNA-seq analysis were generated from Cre^Tg/0^ Ppp2r2a^L/L^ x Cre^0/0^ Ppp2r2a^L/L^ breeding pairs. All aspects of animal care and experimentation were approved by The Alfred Research Alliance Animal Ethics Committee. Mice were housed in a temperature-controlled environment with a 12-hour light/dark cycle and fed standard chow *ad libitum*.

**Figure 1:**
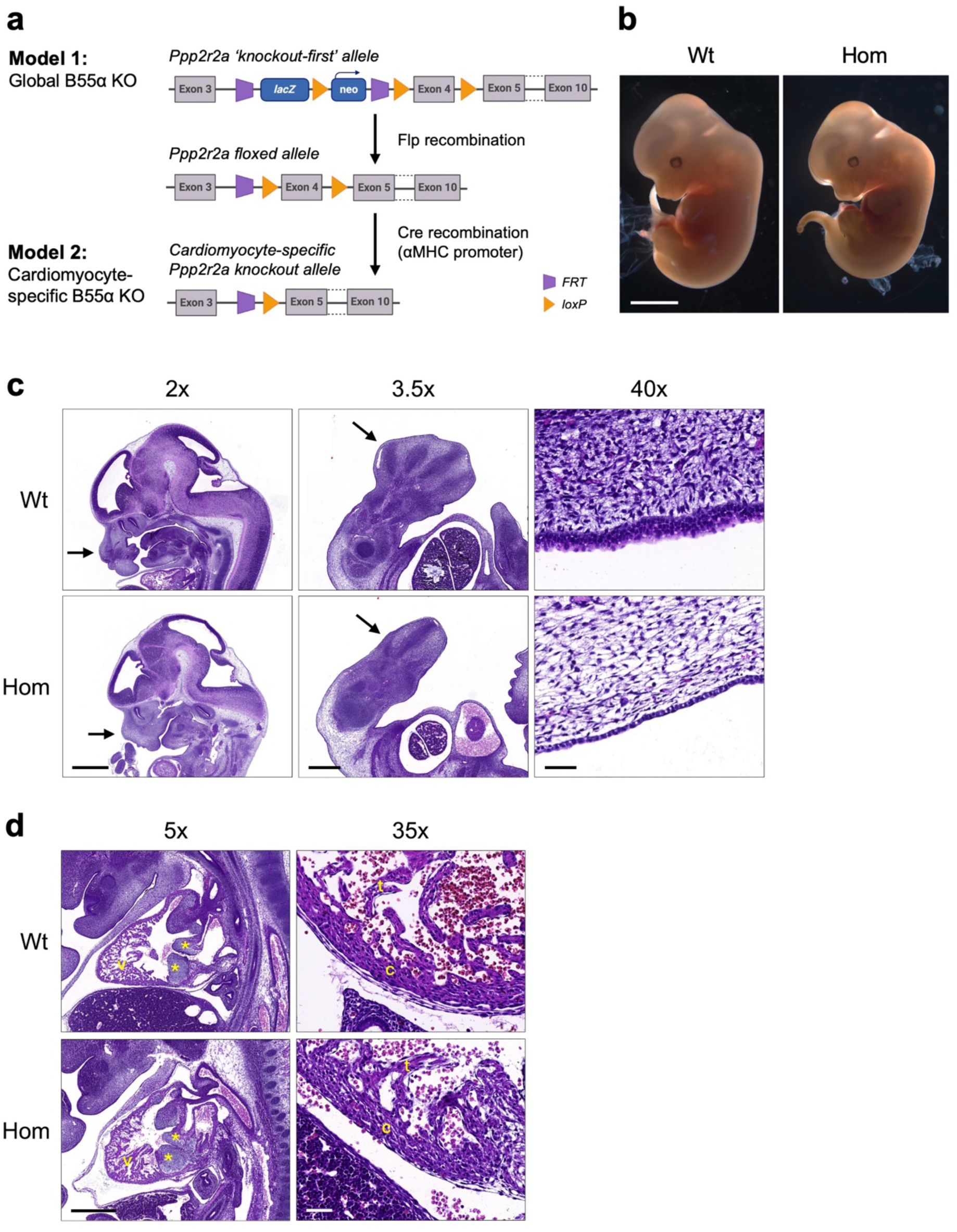
Homozygous knockout of B55α causes embryonic lethality without overt cardiac defects. a) Schematic of the *Ppp2r2a* ‘knockout-first’ allele used to produce B55α heterozygous (Het) and homozygous (Hom) knockout mice (Model 1: Global B55a KO). Floxed *Ppp2r2a* mice were produced by removing the lacZ/neo cassette with Flp recombinase. Cardiomyocyte-specific B55α knockout mice (Model 2) were produced by removing exon 4 with Cre recombinase under control of the αMHC promoter. b) Whole mount images of wildtype (Wt) and Hom littermates at 12.5 dpc, imaged using an AxioCamMRc5 camera with a 1x objective. Scale bar 2 mm. c) Images of Wt and Hom embryos at 12.5 dpc, stained with haematoxylin & eosin and imaged using a Pannoramic Scan II slide scanner fitted with a 20x/NA 0.8 objective and a CCD camera. Left panel shows head (scale bar 1 mm), middle panel shows hindlimb (scale bar 0.5 mm), right panel shows skin layer (scale bar 0.05 mm). d) Images of Wt and Hom hearts at 12.5 dpc, stained with haematoxylin & eosin and imaged using a Pannoramic Scan II slide scanner fitted with a 20x/NA 0.8 objective and a CCD camera. Left panel shows heart cross-section (scale bar 0.5 mm), right panel shows ventricular wall (scale bar 0.05 mm).

### 3.2. Timed matings for collection of embryos at 12.5 dpc

Het breeding pairs were mated. Pregnancy was identified via the presence of a vaginal plug, and midday was designated as 0.5 days post coitum (dpc). Pregnant females were anesthetised with pentobarbital (∼30 mg intraperitoneal) and euthanised via cervical dislocation. An incision was made in the lower abdomen, and the uterine horn removed and immediately placed on sterile gauze in an ice slurry. Embryos were carefully dissected under microscope guidance and placed in cold 4% paraformaldehyde (PFA) with gentle shaking overnight.

### 3.3. Embryo histology

Embryos were dehydrated, paraffin-embedded in a sagittal orientation and 5 µm sections obtained at the level of the heart. Sections were stained with hematoxylin & eosin (H&E) and imaged using a Pannoramic Scan II (3D Histech, Budapest, Hungary), using a Carl Zeiss Plan-Apochromat 20x/NA 0.8 objective and a Point Grey Grasshopper 3 CCD monochrome camera. Digital slide images were viewed using SlideViewer software version 2.6 (3D Histech, Budapest, Hungary). Embedding and processing was conducted by the University of Melbourne Histology Platform (Melbourne, Australia) and slide scanning performed by Phenomics Australia (Melbourne, Australia). Additionally, four paraffin-embedded embryos (3 Hom, 1 Wt) were submitted to Phenomics Australia for histopathological evaluation^29–34^ by two independent pathologists.

### 3.4. Echocardiography

Echocardiography was performed on anaesthetised mice (1.8% isoflurane) using either an iE33 ultrasound machine with a 15-megahertz (mHz) linear array transducer (Phillips, Amsterdam, Netherlands) or a Vevo 2100 High Frequency Ultrasound System with a MS550D transducer (Visual Sonics, Toronto, ON, Canada). Short axis M-mode images were used to measure interventricular septum (IVS) and left ventricular posterior wall (LVPW) thicknesses at diastole, as well as internal dimensions at diastole (LVID; d) and systole (LVID; s). Dimensions were measured from three beats per echocardiogram and averaged. Fractional shortening and LV mass were calculated using the following equations, respectively: [(LVID; d – LVID; s)/LVID; d] x 100%, [(IVS + LVPW + LVID; d)^3^ – LVID; d ^3^]. Investigators were blinded to genotype during image acquisition and analysis. All analyses were independently validated by the Baker Heart & Diabetes Institute Preclinical Microsurgery & Imaging Platform (Melbourne, Australia).^35^

### 3.5. Tissue collection

Adult mice were anesthetised with pentobarbital (∼30 mg intraperitoneal) and euthanised via cervical dislocation. The heart was quickly removed, washed in cold PBS, excess moisture removed with sterile gauze, and weighed. The atria were dissected and weighed. Ventricles were cut along the short axis, separating the base and apex. The base was fixed in cold 4% PFA for processing and paraffin embedding. The apex was divided evenly along the long axis and snap-frozen in liquid nitrogen for protein and RNA extraction. Other tissues including lungs, liver, kidney, and spleen were weighed and snap-frozen in liquid nitrogen.

### 3.6. Gene expression by quantitative PCR

Total RNA was extracted from ventricles using TRI-Reagent (Sigma Aldrich, St. Louis, MO, US), according to the manufacturer’s instructions. Nanodrop spectrophotometry (Thermo Scientific, Waltham, MA) was used to quantify RNA. A High-Capacity RNA-to-cDNA kit (Thermo Scientific, Waltham, MA) was used to reverse transcribe 2 µg RNA to cDNA according to the manufacturer’s instructions. RT-qPCR was conducted using Taqman gene expression assays and Taqman Fast Universal Master Mix (Thermo Fisher Scientific, Waltham, MA) or primers and SYBR Green PCR Master Mix (Thermo Fisher Scientific, Waltham, MA). Targets were amplified using a Quant Studio ViiA7 Real-Time PCR System. The 2^−ΔΔCt^ method was used for calculation of fold difference using hypoxanthine phosphoribosyltransferase 1 (*Hprt1*) as a reference gene. For SYBR Green reactions, primer efficiencies were calculated according to the following equation: E=10^[-1/slope]^, using the slope of a standard curve generated using serial dilutions of cDNA. Melt curves were performed to assess specificity of the amplification product. Samples were run in triplicate, and samples with replicates with a standard deviation exceeding 0.5 were excluded. Taqman gene expression assays from Thermo Fisher Scientific (Waltham, MA) were as follows: *Nppa* Mm01255747_g1, *Nppb* Mm01255770_g1, *Myh6* Mm00440359, *Myh7* Mm01319006, *Col1a1* Mm00801666_g1*, Col3a1* Mm00802300_m1*, Ctgf* Mm01192932_g1*, Hprt1* Mm01545399_m1. SYBR Green primer sequences were as follows: *Ppp2r2a* fwd 5’-CCGTGGAGACATACCAGGTA-3’, rev 5’-AACACTGTCAGACCCATTCC-3’; *Ppp2r2c* fwd 5’-AGCGGGAACCAGAGAGTAAG-3’, rev 5’-GTAGTCAAACTCCGGCTCG-3’; *Ppp2r2d* fwd 5’-TTACGGCACTACGGGTTCCA-3’, rev 5’-TTCGTCGTGGACTTGCTTCT-3’; *Hprt1* fwd 5’-TCCTCCTCAGACCGCTTTT-3’, rev 5’-CCTGGTTCATCATCGCTAATC’-3’.

### 3.7. Protein analysis

Ventricles were homogenised in lysis buffer [20 mM Tris-HCl (pH 7.4), 127 mM NaCl, 10% glycerol, 1% IGEPAL, 20 mM NaF, 10 mM EGTA, 1 mM sodium pyrophosphate, 1 mM vanadate, 1 mM PMSF, 4 µg/mL pepstatin, 4 µg/mL aprotinin, 4 µg/mL leupeptin], incubated on ice for 15 min, and centrifuged at 4°C for 15 min at 16,000 *g* to pellet cellular debris. Protein lysates (80 µg) and molecular weight marker (Precision Plus Protein All Blue Prestained Protein Standard, Bio-Rad, 1610373) were separated using 7.5% or 10% SDS-PAGE gels, transferred to PVDF membranes, and incubated with primary antibody overnight at 4°C or 1.5 hours at room temperature for HRP-conjugated secondary antibodies. Chemiluminescent signals were quantified using ImageJ pixel analysis (US National Institutes of Health) or Genetools analysis software (Syngene, Cambridge, UK). Data were expressed as fold change relative to the control group. Antibodies were as follows: B55α (#sc-81606, Santa Cruz, 1:1000), FoxO1 (#2880, Cell Signaling Technology, 1:2000), pSer256 FoxO1 (#9461, Cell Signaling Technology, 1:1000), pThr24/Thr32 FoxO1/O3a (#9464, Cell Signaling Technology, 1:2000), HDAC5 (#20458, Cell Signaling Technology, 1:500), pSer259 HDAC5 (#3443, Cell Signaling Technology, 1:500), PP2A-C (#2038, Cell Signaling Technology, 1:1000), PP2A-A (#2039, Cell Signaling Technology, 1:250), α-tubulin (#2144, Cell Signaling Technology, 1:2500) and vinculin (#V9131, Sigma Aldrich, 1:5000). All uncropped blots are included in the Data Supplement.

### 3.8. RNA-sequencing

Left ventricle tissue (∼20 mg) was mechanically homogenised in TRI reagent with a 5 mm steel bead at 30 Hz for 2 x 30 s (TissueLyser II, Qiagen) then centrifuged for 10 min at 10,000 *g*. RNA was extracted from the supernatant using spin columns with DNase-I treatment (Zymo DirectZol RNA Miniprep kit #R2050, Integrated Sciences, Australia). RNA concentration and purity was determined spectrophotometrically (Nanodrop, Thermo Fisher, Australia) and RNA integrity was determined by electrophoresis (Tapestation, Agilent). RNA integrity (RIN) values for all samples were ≥7.7. Stranded libraries were prepared from 500 ng total RNA input with poly-A+ selection (TruSeq Stranded mRNA, Illumina). Libraries size and concentration were confirmed prior to sequencing (TapeStation, Agilent). At least 40 million 150 bp paired-end reads per library were generated on the Illumina NovaSeq 6000 platform (Australian Genome Research Facility, Melbourne, Australia). Raw read preprocessing was conducted on the Galaxy Australia platform.^36^ Reads underwent initial quality check with FastQC v0.74, then quality filtering and adapter trimming was conducted with Cutadapt v4.6. Filtered reads were aligned to the mouse reference genome (Mus musculus, Ensembl version GRCm39.104) using STAR v2.7.11a^37^ in 2-pass mode. Gene expression was quantified at the gene level using HTseq-counts v2.0.5 and collated into a counts matrix. Analysis of differential expression was performed using on genes with ≥1 count per million (CPM) in all samples in DESeq2 in iDEP v2.0.^38^ False discovery rate (FDR) was calculated using Benjamini-Hochberg method and genes with an FDR<0.05 were considered differentially expressed (DE). DE genes were analysed for pathway enrichment using Gene Ontology (GO) genesets in iDEP v2.0.

### 3.9. Fibrosis analysis

Paraffin-embedded left ventricles were cut (6 μm sections), stained with picrosirius red and imaged using a Pannoramic Scan II (3D Histech, Hungary) using a Carl Zeiss Plan-Apochromat 20x/NA 0.8 objective and a Point Grey Grasshopper 3 CCD monochrome camera. Images of scanned sections were obtained at 20x magnification (10 regions/heart) using SlideViewer software version 2.6 (3D Histech, Budapest, Hungary). The number of red pixels (collagen) was counted using ImageJ (NIH, Bethesda, USA) and divided by tissue area to calculate the percentage of fibrosis. Analysis was undertaken blinded to sex and genotype.

### 3.10. Data presentation and statistical analysis

Data are presented as scatterplots with bars and lines indicating the mean +/- SEM (for qPCR data, as per Thermo Fisher Scientific’s *Guide to Performing Relative Quantitation of Gene Expression Using Real-Time Quantitative PCR*, 2008 edition) or median +/- IQR (all other data). Fold difference (qPCR and Western blot data) was calculated by dividing by the mean or median of the control group, as appropriate. Gene expression data were analysed using two-sided unpaired *t*-tests (2 groups) or two-way ANOVA followed by Tukey’s or Sidak’s post-hoc tests, as appropriate. All other data were analysed using Mann-Whitney U tests (2 groups) or Kruskal-Wallis tests (6 groups). A *P* value <0.05 was deemed statistically significant. Animal numbers and data exclusions are reported in **Supp Fig 1**.

### 3.11. Data availability

RNA-sequencing datasets from this study have been deposited at the NCBI Gene Expression Omnibus (accession number GSE266131).

## 4. Results

To determine if PP2A-B55α regulates heart size or morphology *in vivo*, we first generated mice with heterozygous (Het) or homozygous (Hom) global disruption of *Ppp2r2a* (Model 1, **Fig 1a**). The knockout-first allele has conditional potential due to the presence of *FRT* and *loxP* sites, allowing removal of the *lacZ* reporter and neomycin selection cassette to generate floxed *Ppp2r2a* alleles for cardiomyocyte-specific Cre-mediated deletion of exon 4 (Model 2, **Fig 1a**).

### 4.1. Homozygous knockout of B55a causes embryonic lethality without overt cardiac defects

The expected Mendelian ratio of Wt:Het:Hom mice in the global knockout mouse strain (Model 1) was 1:2:1. However, no Hom offspring were born to Het breeding pairs, indicating lethality and resorption of Hom embryos *in utero*. Consistent with previous reports in different B55α knockout mouse strains,^11,26^ Hom embryos were smaller than Wt littermates at 12.5 dpc (**Fig 1b**) and started dying between 12.5 and 14.5 dpc (26% Hom at 12.5 dpc, 6% Hom at 14.5 dpc). Histopathological evaluation at 12.5 dpc identified several abnormalities in Hom embryos, including arrested oral/nasal cavity development (**Fig 1c**, left panel), arrested limb bud development (**Fig 1c**, middle panel), and deficient skin layers (**Fig 1c**, right panel). One of the three Hom embryos evaluated also had a smaller liver and displayed hepatocyte necrosis. Hearts of Hom embryos were assessed for the presence of congenital cardiac abnormalities, including atrial or ventricular septal defects, abnormal leaflet formation or myocardial non-compaction, however there were no obvious micromorphological differences compared with Wt (**Fig 1d**).

### 4.2. Reduced expression of B55α did not lead to left ventricular hypertrophy or dysfunction in young mice but induced LV wall thinning in male mice with age

Ventricular B55α protein levels were ∼45% lower in the male Het group vs Wt and ∼70% lower in the female Het group vs Wt (**Fig 2a, b**), allowing us to investigate the impact of reduced B55α expression on cardiac morphology and function in adult mice. There were no differences in the relative abundance of B55α between male and female control hearts (**Supp Fig 2**). We assessed mice at 10-12 weeks of age (representing early adulthood) and at 12 months (representing middle age). By echocardiography, there were no differences in left ventricular wall thicknesses or internal dimensions between Wt and Het mice in the 10-12-week old cohort (**Supp Table 1**). Heart rate and fractional shortening, a measure of left ventricular contractility, were also not different between Wt and Het mice (**Supp Table 1**). At 12 months of age, male Het mice had thinner LV walls compared to Wt mice based on LVPW and IVS thicknesses (**Fig 2c**). This phenotype was less apparent in females, with lower IVS thickness in Het mice but no difference in LVPW (**Fig 2d**). No other differences in echocardiographic parameters were observed (**Fig 2c, d**; **Supp Table 2**).

**Figure 2:**
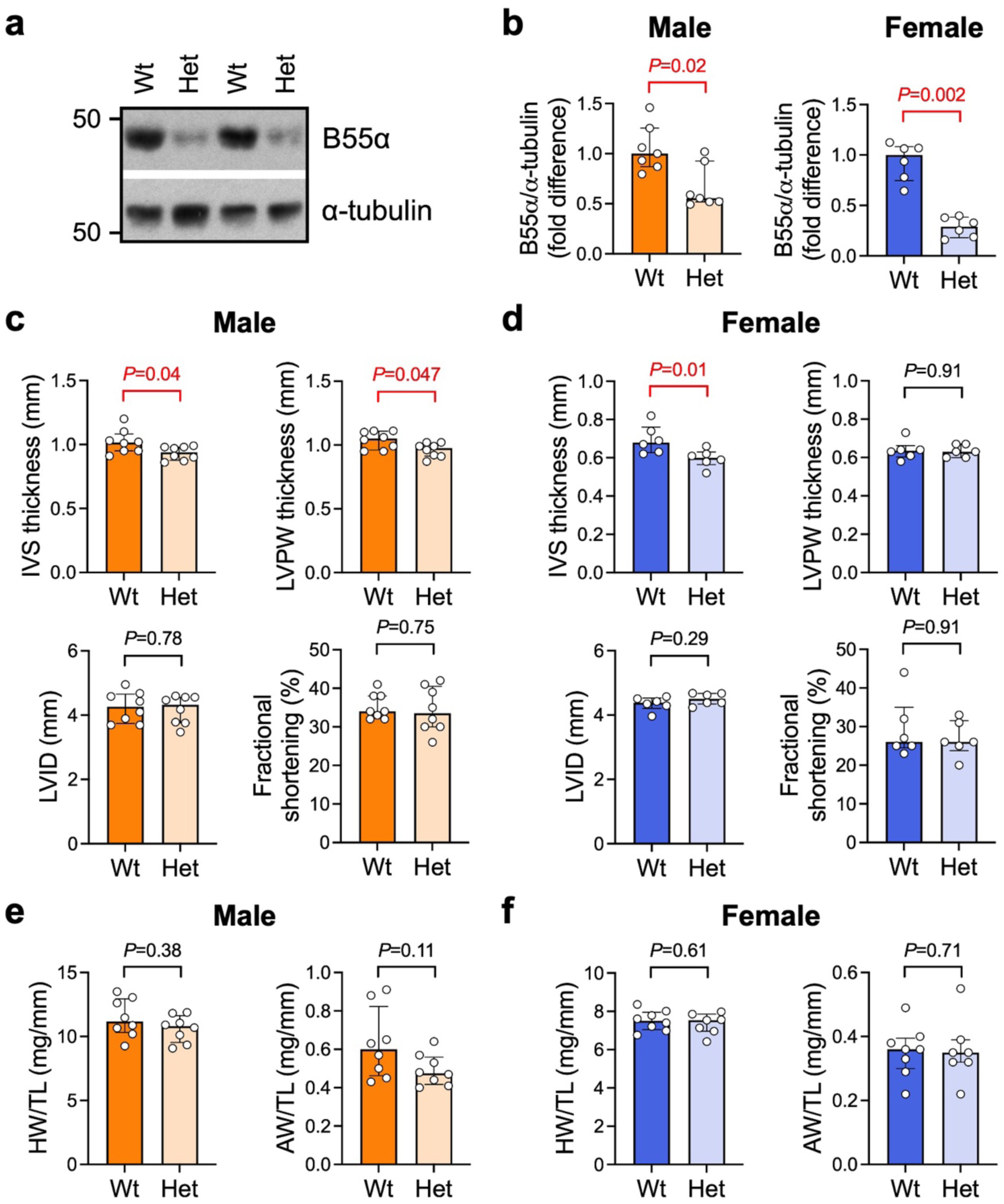
Heterozygous expression of B55α leads to left ventricular wall thinning in male mice at 12 months of age. a) Representative Western blot and b) quantitation of B55α protein expression. Ventricular tissue from 12-month old male and female *Ppp2r2a* wildtype (Wt) and heterozygous (Het) mice at 12 months of age. α-tubulin was used as a loading control. Data are expressed as fold difference vs Wt. Bars show median +/- IQR. Mann-Whitney U tests. *n*=6-7 per group. c, d) Interventricular septum (IVS) thickness, left ventricular posterior wall (LVPW) thickness, LV internal dimension (LVID) and fractional shortening, assessed by M-mode echocardiography in anaesthetised mice at 12 months of age. Bars show median +/- IQR. Mann-Whitney U tests. *n*=6-8/group. e, f) Heart weight (HW) and atria weight (AW) normalised to tibia length (TL) of mice at 12 months of age. *n*=6-8/group. Bars show median +/- IQR. Mann-Whitney U tests.

Assessment of organ weights at 10-12 weeks and at 12 months of age revealed no differences in heart weight or atria when normalised to body weight or tibia length, although atria tended to be smaller in male Het vs Wt in the younger cohort (**Fig 2e, f**; **Supp Table 3, Supp Table 4**). Lung weight was within the normal range, consistent with the absence of cardiac pathology (which can lead to pulmonary congestion; **Supp Table 3**, **Supp Table 4**). Liver weight was lower in female Het vs Wt in the younger cohort (**Supp Table 3**) and tended to be higher in male Het vs Wt in the older cohort (see **Supp Table 4**). Variations in the liver were also observed in Hom embryos at 12.5 dpc (**Section 4.1**), indicating a possible hepatic function for B55α. We did not measure blood pressure in these mice, which is a limitation of the study.

To investigate if the mild LV wall thinning observed in Het mice at 12 months of age was accompanied by differences in the expression of genes involved in pathological cardiac remodelling, we performed qPCR on ventricular lysates (**Supp Fig 3a, b**). Levels of cardiac stress markers atrial and B-type natriuretic peptides (ANP and BNP, encoded by *Nppa* and *Nppb* respectively) did not differ in Het mice compared to Wt in either sex. There were no differences in the expression of myosin heavy chain isoforms αMHC or βMHC (encoded by *Myh6* and *Myh7* respectively). Expression of type I and III collagens (*Col1a1* and *Col3a1*) and connective tissue growth factor (*Ctgf*) were comparable between genotypes, suggesting an absence of cardiac fibrosis. We also assessed the expression of blood and lymphatic vessel markers *(Cd31* and *Lyve1*, respectively) as impaired angiogenesis or lymphangiogenesis in the heart is deleterious,^39,40^ and a previous study reported vascular and lymphatic vessel defects in skin of embryos lacking B55α.^11^ We observed a significant reduction in mRNA, but not protein, levels of the lymphatic vessel marker *Lyve1* in male Het ventricles vs Wt (**Supp Fig 3c, d**), while *Cd31* expression was not different between genotypes (**Supp Fig 3c**). *Lyve1* mRNA and protein levels were not different in female Het and Wt hearts (**Supp Fig 3e, f**). *Cd31* mRNA expression tended to be lower in female Het mice vs Wt, however the effect was small (∼10% decrease, *P*=0.05, **Supp Fig 3e**).

### 4.3. Loss of B55α in cardiac myocytes does not impact postnatal cardiac growth, but alters the transcriptional proWile in male mice

To circumvent the effects of B55α deletion on embryonic development and investigate the role of cardiomyocyte B55α in postnatal heart growth, we next characterised the phenotype of mice with cardiomyocyte-specific knockout (cKO) of B55α using an αMHC-Cre line (Model 2, **Fig 1a**). We confirmed loss of B55α in ventricular tissue of 10-12-week old cKO mice by Western blotting. As expected, B55α protein levels were significantly lower (∼90% decrease in expression) in both male and female cKO hearts compared to floxed controls (FC, **Fig 3a-d**). We also examined the expression of the other components of the heterotrimeric PP2A holoenzyme. We observed a significant decrease (∼35%, *P*=0.03) in PP2A-A protein levels in female cKO hearts, but no differences in the expression of PP2A-C (**Fig 3a-d**). Deletion of *Ppp2r2a* in the heart was not associated with differences in the expression of other PP2A B55 subunit isoforms, i.e. *Ppp2r2c* (encoding B55γ) or *Ppp2r2d* (encoding B55δ; **Fig 3e, f**). We also observed no differences in HDAC5 phosphorylation at Ser259 between genotypes in either sex (**Fig 3g, h**), suggesting that HDAC5 is not phospho-regulated by PP2A-B55α in the heart under normal physiological conditions.

**Figure 3:**
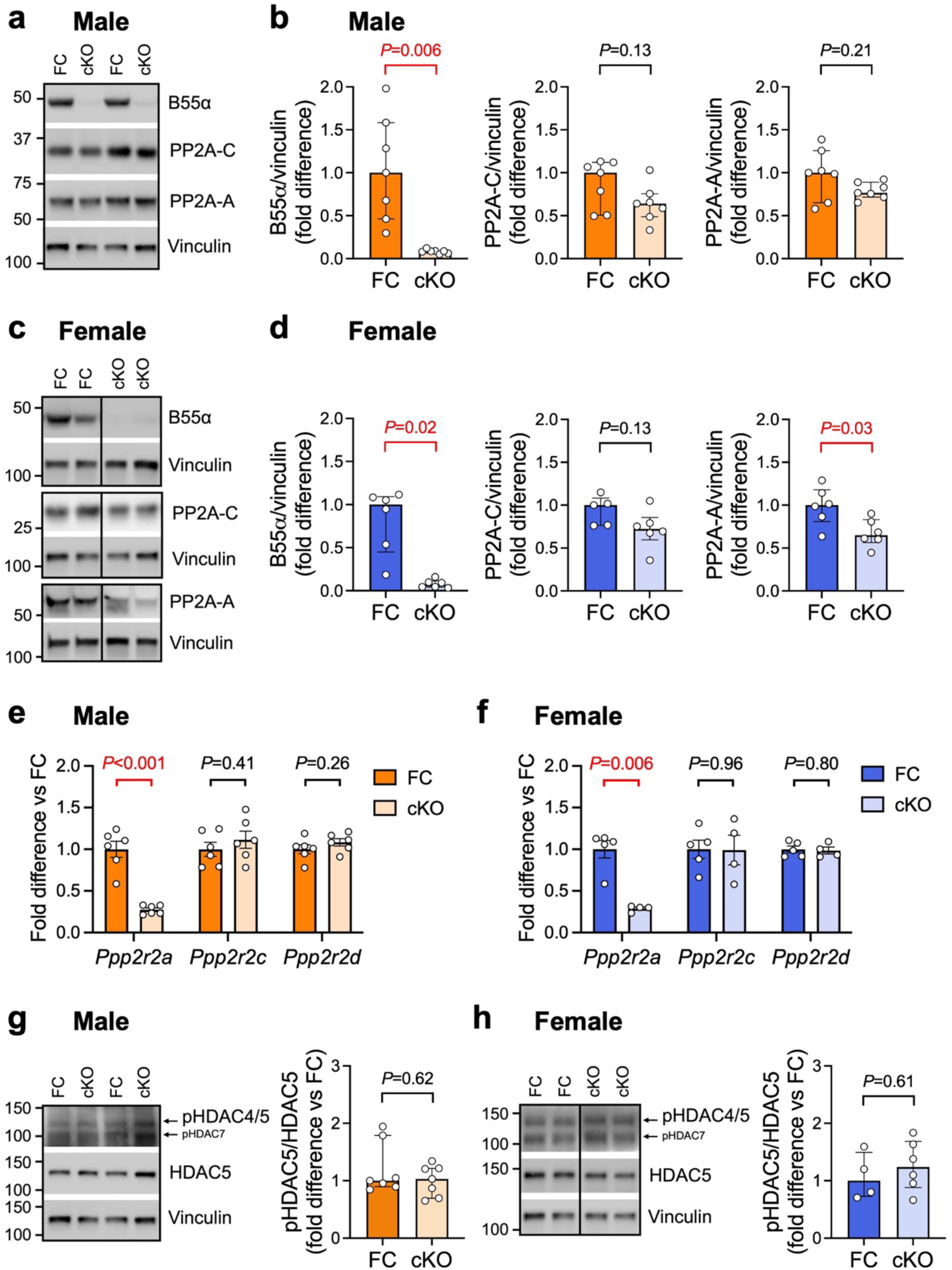
Effects of cardiomyocyte-specific deletion of *Ppp2r2a* on PP2A subunit expression. a, c) Representative Western blots and b, d) quantitation of B55α, PP2A catalytic (PP2A-C) and scaffolding (PP2A-A) subunits in ventricular tissue from 10-12 weeks old cardiomyocyte-specific B55α knockout (cKO) mice and floxed controls (FC). Vinculin was used as a loading control. Data are expressed as fold difference vs FC. Bars show median +/- IQR. Mann-Whitney U tests. *n*=6 per group for all analyses except PP2A-C (*n*=5 for female FC). Vertical lines indicate where images of Western blots have been spliced so the relevant genotypes could be displayed side by side. e, f) Quantitative PCR analysis of PP2A B55 subunits *Ppp2r2a* (encoding B55α), *Ppp2r2c* (encoding B55γ) and *Ppp2r2d* (encoding B55δ). Data are expressed as fold difference relative to the FC group. Bars show mean +/- SEM. Unpaired t-tests. *n*=4-6/group. g, h) Representative Western blots and quantitation of HDAC5 phosphorylation at Ser259 relative to total HDAC5. Data are expressed as fold difference vs FC. Bars show median +/- IQR. Mann-Whitney U tests. *n*=4-7 per group.

At 10-12 weeks of age, there were no detectable differences in LV wall thicknesses or chamber dimensions between any of the genotypes (**Fig 4a, b**; **Supp Table 5**). Fractional shortening tended to be lower in male cKO vs FC, but the effect was small (∼10% decrease, *P*=0.10). Heart, atria and lung weight of both male and female cKO mice were comparable with controls (**Fig 4c, d**; **Supp Table 6**), indicating the absence of pathological cardiac hypertrophy and dysfunction.

**Figure 4:**
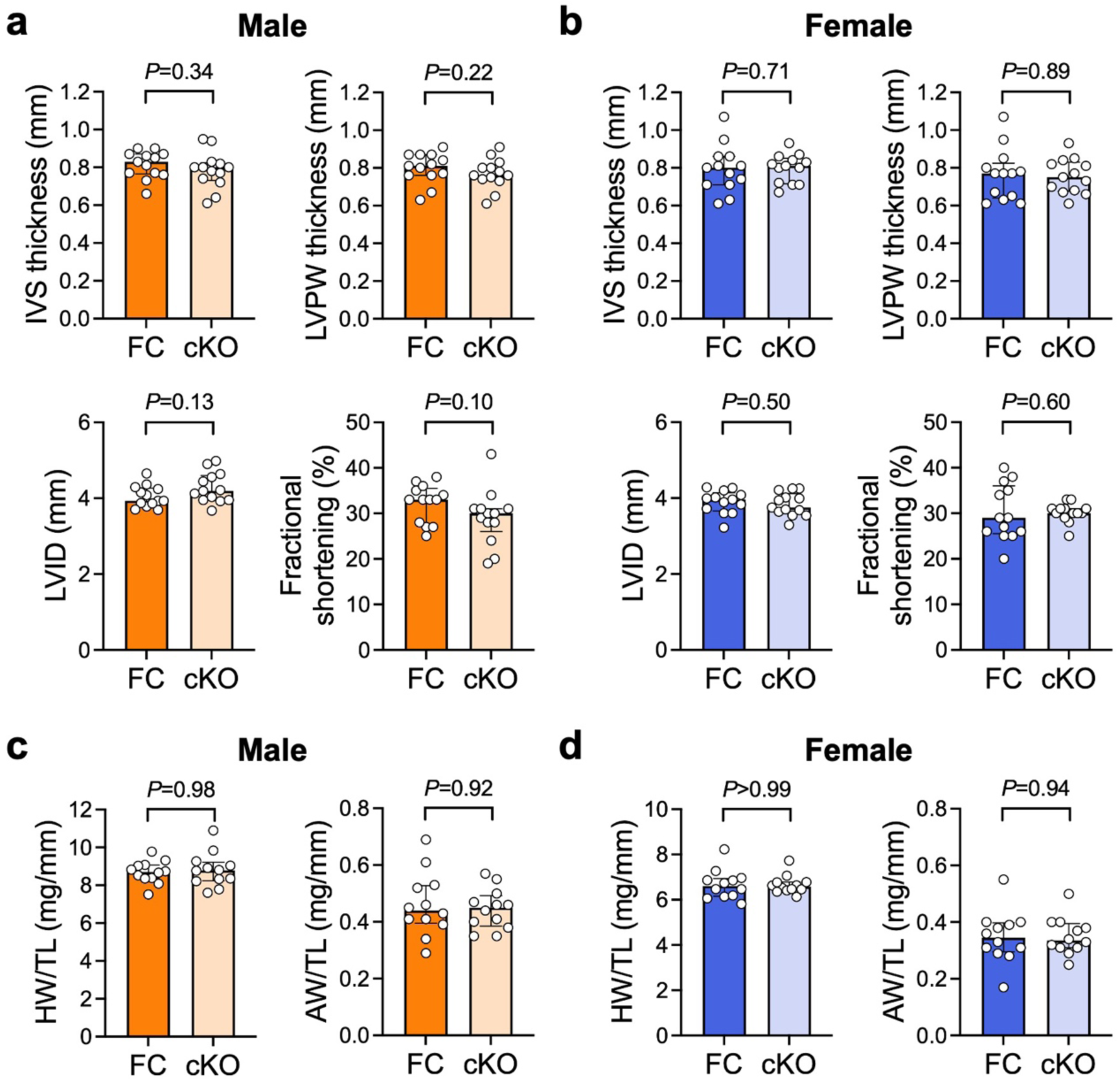
Left ventricular structure and function was normal in mice with cardiomyocyte-specific knockout of B55α at 10-12 weeks of age. a) Interventricular septum (IVS) thickness, left ventricular posterior wall (LVPW) thickness, LV internal dimension (LVID) and fractional shortening in 10-12 week old cardiomyocyte-specific B55α knockout (cKO) mice and floxed controls (FC), quantified from M-mode echocardiograms. Lines indicate the median. Mann-Whitney U tests. *n*=13/group. b) Heart weight (HW) and atria weight (AW) normalised to tibia length (TL) in FC and cKO mice at 10-12 weeks of age. Lines indicate the median. Mann-Whitney U tests. *n*=12/group.

Interestingly, we observed differences in the expression of genes involved in extracellular matrix and myofilament composition in ventricular tissue of male mice (**Fig 5a**). Expression of type I collagen (*Col1a1*) and connective tissue growth factor (*Ctgf*) were modestly increased in male cKO hearts compared with FC (∼30% increase in *Col1a1*, *P*=0.04; ∼70% increase in *Ctgf*, *P*=0.02, **Fig 5a**). Type III collagen expression also tended to be higher (∼20% increase in *Col3a1*, *P*=0.08, **Fig 5a**). *Myh7* (encoding βMHC) was also significantly higher in male cKO hearts vs FC (∼70% increase, *P*=0.02, **Fig 5a**). Upregulation of the foetal βMHC isoform is frequently observed in cardiac disease settings.^16,41,42^ Coupled with our observation of increased lower fractional shortening in male cKO mice vs FC, this gene expression profile could indicate the early onset of pathology in male cKO mice, although increased expression of the βMHC isoform is typically accompanied by a reciprocal decrease in the expression of the αMHC isoform.

**Figure 5:**
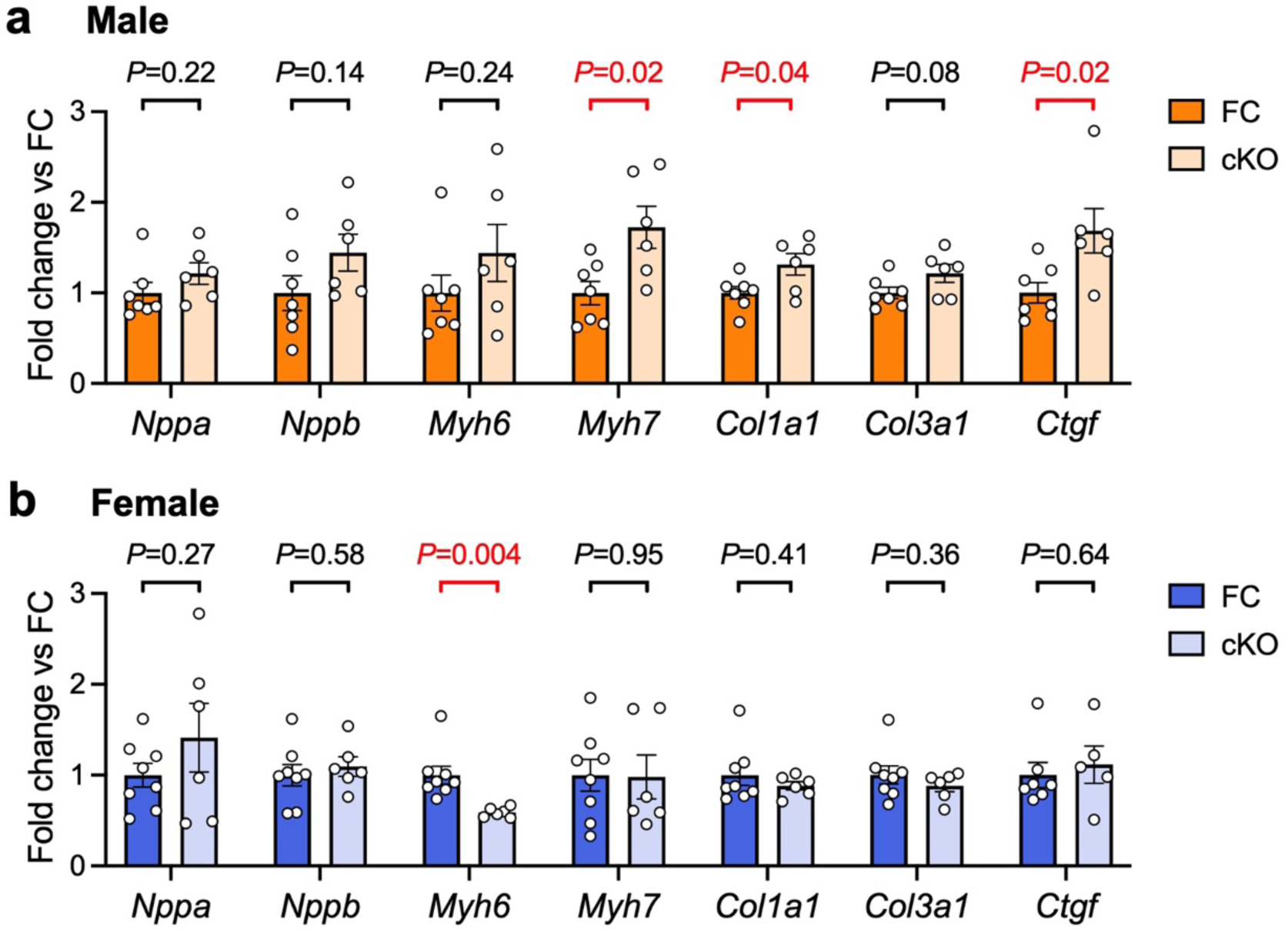
Loss of cardiomyocyte B55α alters the expression of myosin heavy chain isoforms and pro-Qibrotic genes. a, b) Quantitative PCR analysis of genes that are typically dysregulated in settings of pathological cardiac hypertrophy and fibrosis. Ventricular tissue from male and female cardiomyocyte-specific B55α knockout (cKO) and floxed control (FC) mice. Data are expressed as fold difference relative to the FC group. Bars show mean +/- SEM. Unpaired t-tests. *n=*6-8 per group for all analyses except *Ctgf* (*n*=5 for female cKO).

We and others have previously shown that the αMHC-Cre transgenic mouse strain used in this study does not have increased collagen expression or fibrosis at 6-8 months of age.^20,28^ However, as another αMHC-Cre strain (with much higher levels of Cre expression) develop cardiac fibrosis,^28^ we compared *Col1a1*, *Col3a1* and *Ctgf* expression in hearts of male FC, cKO and Cre control mice. Our analyses revealed no differences in expression between the FC and Cre control groups (**Supp Fig 4**), confirming that the modest increases in expression observed in cKO hearts were due to deletion of B55α and not an off-target effect of Cre expression.

In female cKO mice, increases in fibrotic gene expression were not present (**Fig 5b**). We observed reduced expression of *Myh6*, a known MEF2 transcriptional target encoding the dominant adult αMHC isoform (∼40% decrease, *P*=0.004, **Fig 5b**). Expression of the natriuretic peptide genes *Nppa* and *Nppb* were not different in male or female cKO hearts compared to control.

To further investigate differences in cardiac gene expression arising from loss of cardiomyocyte B55α, we performed RNA-seq analysis on a separate cohort of FC and cKO mice. Interestingly, and consistent with the findings of our qPCR analysis (**Fig 5a, b**), we observed a greater number of genes that were differentially expressed due to B55α deletion in the male heart compared with the female heart (**Fig 6a, b; Supp Table 7**). Remarkably, only 5 genes (in addition to *Ppp2r2a*) were affected by B55α deletion in the female heart (**Fig 6b**), while 543 were differentially expressed in male cKO vs FC (FDR<0.05, **Fig 6a**). *Eya1* was the only gene that was significantly upregulated in both male and female hearts with B55α deletion (**Fig 6a, b**). Eya family members are transcriptional co-activators that have been shown to physically interact with B55α, directing PP2A activity towards transcription factors such as c-Myc.^43^ The physiological significance of this interaction has not been investigated in the heart.

**Figure 6:**
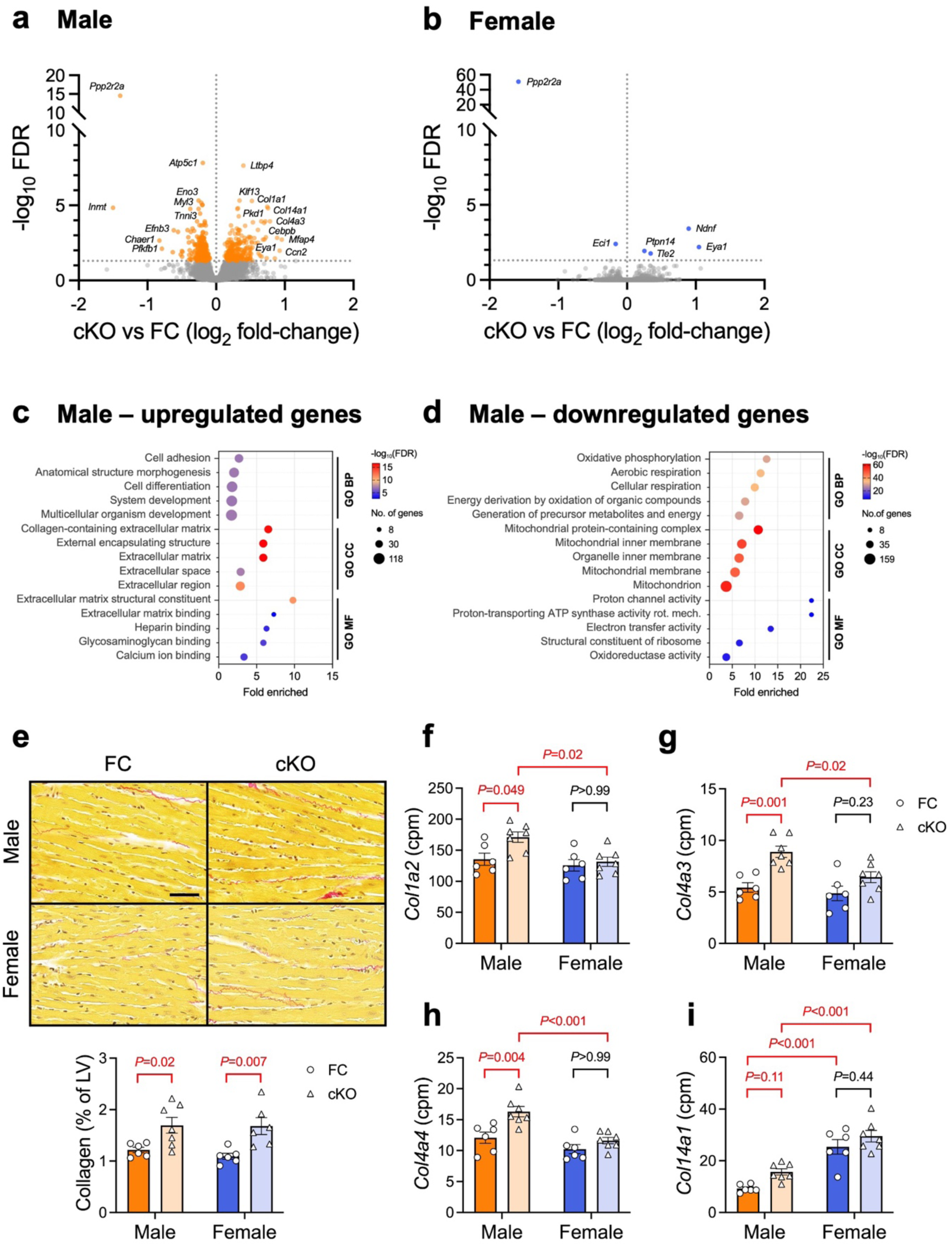
Loss of cardiomyocyte B55α had a greater impact on cardiac gene expression in males compared with females. a, b) Volcano plots showing genes that were differentially expressed between cKO and FC hearts. RNA-seq analysis of ventricular tissue from male and female cardiomyocyte-specific B55α knockout (cKO) and floxed control (FC) mice. FDR<0.05, *n=*6-7/group. c, d) Pathway enrichment analysis showing Gene Ontology terms that were over-represented in the set of genes that were significantly upregulated or downregulated in male cKO hearts vs FC. *n=*6-7/group. e) Representative LV sections stained with picrosirius red and quantitation of collagen (expressed as a percentage of left ventricular area). Scale bar: 50 μm. Two-way ANOVA followed by Sidak’s posthoc tests. *n*=6-7/group. f-i) Relative mRNA abundance (in counts per million, cpm) of collagen isoforms in FC (circles) and cKO (triangles) hearts. Two-way ANOVA followed by Tukey’s posthoc tests. *n*=6-7/group.

Pathway enrichment analysis of genes that were upregulated in male cKO vs FC hearts revealed an over-representation of Gene Ontology terms associated with cell/tissue/organism development and morphogenesis (Biological Process), extracellular matrix (Cellular Component), and ECM structural constituents and binding (Molecular Function, **Fig 6c; Supp Table 8**). Amongst genes that were downregulated in male cKO vs FC hearts, there was an over-representation of terms associated with oxidative phosphorylation and energy production (Biological Process), mitochondria (Cellular Component), and activity of the electron transport chain (Molecular Function, **Fig 6d**). These data suggest a role for B55α in the maintenance of ECM composition and mitochondrial energy metabolism in the male heart.

Analysis of LV cross-sections stained with picrosirius red revealed that upregulation of ECM-associated genes in male cKO mice translated to a small increase in collagen protein deposition (**Fig 6e**). Surprisingly, we detected a similar increase in LV collagen content in female cKO mice vs FC, despite no differences in the expression of *Col1a1* or *Col3a1* (**Fig 5b**, **Fig 6b**). Examination of other collagen species quantified by RNA-seq revealed a significant effect of genotype and sex for *Col1a2*, *Col4a3*, *Col4a4* and *Col14a1* (*P*<0.05 for all, **Fig 6f-i**). *Col4a3* and *Col4a4* expression tended to be higher in female cKO hearts vs FC, although these differences were not as pronounced as in male hearts and were not statistically significant (**Fig 6g-h**). *Col14a1* expression was significantly higher in female hearts compared with male hearts (∼2.7-fold increase in female FC vs male FC, *P*<0.001, **Fig 6i**). Collectively, these data identify cardiomyocyte B55α as a regulator of the cardiac ECM.

In addition, we investigated the phosphorylation status of FoxO1, a reported target of B55α (at Ser256 and Thr24)^18^ and key transcription factor regulating mitochondrial homeostasis and cardiac energy metabolism.^44,45^ FoxO1-dependent gene transcription is regulated by phosphorylation, with Ser256 dephosphorylation promoting the nuclear accumulation of FoxO1 and Ser256 phosphorylation promoting cytosolic accumulation in cardiomyocytes.^46^ We detected no differences in the phosphorylation of FoxO1 at Ser256 or Thr24 between genotypes (**Supp Fig 5**), suggesting that FoxO1 is unlikely to have contributed to the gene expression differences we observed in hearts of B55α cKO mice.

## 5. Discussion

The aim of this study was to determine if the PP2A regulatory subunit B55α is required for heart growth during normal physiological development. We generated mice with global or cardiomyocyte-specific deletion of B55α and examined the cardiac phenotype during embryogenesis, early adulthood (10-12 weeks) and at 12 months of age. The key findings of this study are: 1) B55α is not required for physiological heart growth during embryogenesis or the postnatal period; 2) in male mice, reduced B55α expression leads to mild thinning of the ventricular walls with ageing; 3) cardiomyocyte-specific deletion of B55α in male mice increases the expression of genes involved in extracellular matrix composition and decreases the expression of genes involved in mitochondrial energy production; 4) loss of B55α increases collagen accumulation in both male and female mouse hearts; and, 5) under basal conditions, B55α is not a major regulator of the female cardiac transcriptome. Collectively, these data demonstrate that the heart can maintain a relatively normal growth trajectory and function, even if B55α protein levels are markedly reduced. This may have implications for PP2A-targeting therapeutics (discussed in **Section 5.4**). Our data reveal sex-specific roles for B55α in cardiomyocytes and point to a possible role for B55α signalling in cardiac endothelial cells during ageing.

Our finding of embryonic lethality in Hom embryos is consistent with two previous publications.^11,26^ Mice with CRISPR/Cas9-mediated deletion of *Ppp2r2a* exon 4 die between 10.5 dpc and birth, with epidermal defects visible as early as 10.5 dpc and limb and digit defects evident at later stages.^26^ Mice with PGK-Cre-mediated (i.e. ubiquitous) deletion of exon 6 die between 12.5 and 15.5 dpc, and display reduced blood and lymphatic vessel density in skin.^11^ This was attributed to loss of B55α in endothelial cells, as conditional deletion of B55α in this cell type recapitulated the phenotype.^11^ Neither study examined the cardiac phenotype of B55α knockout embryos.

In the current study, the cardiac phenotype of embryos with homozygous deletion of B55α was assessed at 12.5 dpc, a developmental timepoint when atrial and ventricular septation is occurring and the ventricular walls are thickening due to myocardial cell proliferation.^32^ Evaluation of H&E-stained sections at 50 μm intervals spanning the heart revealed no overt signs of abnormal cardiac development in Hom embryos. It is possible that more detailed investigations could reveal subtle phenotypic changes that are not detectable by H&E staining. Based on the findings of Ehling and colleagues,^11^ it would not be surprising if hearts of knockout mice displayed reduced blood and lymphatic vessel density. Endothelial cells are the most abundant non-myocyte cell type in the heart,^47^ and the development of both the coronary and lymphatic vasculature commences prior to embryonic day 14.5, coinciding with the occurrence of embryonic lethality in our model.^48–50^ In support of a role for B55α in cardiac endothelial cell populations, we observed a decrease in mRNA levels of the lymphatic vessel marker *Lyve1* in hearts of male mice with reduced B55α expression at 12 months of age, although this was not associated with a detectable reduction in LYVE1 protein. In the heart, lymphatic vasculature is critical for the maintenance of fluid equilibrium as well as nutrient and immune cell transport,^49–54^ and *Lyve1* knockout mice had worse cardiac function and more pronounced wall thinning than control mice in response to myocardial infarction.^39^ Thus, reduced B55α in endothelial cell populations may make the heart more susceptible to pathological cardiac remodelling and dysfunction with ageing or in settings of cardiac stress or injury.

After birth, the heart responds to sustained increases in workload by increasing heart mass.^55^ This is achieved via the enlargement of heart muscle cells (cardiomyocyte hypertrophy). The signalling pathways regulating cardiomyocyte hypertrophy are complex, involving numerous receptors and stretch sensors, signalling mediators and transcription factors. A key signalling axis regulating cardiomyocyte hypertrophy in disease settings is the HDAC5/MEF2 axis.^56^ The adult heart expresses MEF2A and MEF2D isoforms, which form homodimers or heterodimers with other transcription factors (e.g. GATA, NFAT and STAT3) and bind to DNA promoter regions to modulate gene transcription.^57–59^ A seminal study from the Olson group showed that MEF2D activation is sufficient and necessary for pathological cardiac remodelling,^16^ while MEF2A may play a protective or a pathological role.^60,61^ The subcellular distribution of HDAC5 is a key determinant of MEF2 transcriptional activity. When localised to the nucleus, HDAC5 represses MEF2. Nuclear export and cytosolic accumulation of HDAC5 following phosphorylation by protein kinases alleviates the repressive interaction with MEF2 and facilitates the transcription of MEF2-dependent genes.^62–67^

We previously showed that B55α targets the PP2A core enzyme to HDAC5 in cardiomyocytes.^12^ We hypothesised that loss of B55α would lead to cardiac hypertrophy, i.e. via hyper-phosphorylation of HDAC5 and activation of MEF2. However, we observed no effect of cardiomyocyte-specific deletion of B55α on LV morphology or heart weight in male or female mice at 10-12 weeks of age. We also observed no differences in the phosphorylation of HDAC5 at Ser259, a key determinant of HDAC5 subcellular localisation which we previously showed is dephosphorylated in a B55α-dependent manner.^12^ It is important to note that our *in vitro* findings were observed in response to β-adrenergic receptor stimulation with isoproterenol, a setting of acute cardiac stress. While B55α was found to interact with HDAC5 in vehicle-treated cardiomyocytes in these experiments, this interaction increased ∼3-fold in response to isoproterenol stimulation and was accompanied by a ∼90% decrease in HDAC5 phosphorylation at Ser259. Our new findings suggest that PP2A-B55α is unlikely to be a major regulator of HDAC5 phosphorylation under basal conditions *in vivo* (i.e. in the absence of β-adrenergic stimulation) and may exert its effects on gene transcription via phospho-regulation of other proteins.

We found that cardiomyocyte-specific deletion of B55α in male hearts was associated with widespread changes in cardiac gene transcription (>500 differentially expressed genes), while the female cardiac transcriptome was largely unresponsive to B55α deletion (<10 differentially expressed genes). These data suggest that the PP2A-B55α holoenzyme may play a more dominant role in the male heart than the female heart under basal conditions, despite comparable B55α protein abundance. PP2A can bind to both the estrogen receptor^68^ and the androgen receptor,^69^ but further work is required to investigate B subunit involvement in PP2A regulation of sex hormones. To date, the role of PP2A in the heart has been studied predominantly in male animals. Sex differences have not been evaluated in a setting of PP2A-C deletion or overexpression.^70–73^ In mice with global deletion of B56α, there were no differences in LV dimensions or HW/TL ratio in males or females assessed at two months of age.^74^ At six months of age, hearts of mice with homozygous knockout of B56α were smaller in males, but not in females, revealing a sex- and age-specific effect of this regulatory B subunit isoform on cardiac morphology.^74^ Further work is required to investigate the relative abundance, subcellular localisation and activity of different PP2A holoenzymes in the male and female heart.

Hearts of male cardiomyocyte-specific B55α knockout mice had increased expression of extracellular matrix-related genes, leading to a small increase in LV collagen content, and decreased expression of genes involved in mitochondrial energy metabolism. Genes associated with the GO Cellular Component term ‘extracellular matrix’ (see **Supp Table 8**) included collagens (*Col1a1, Col1a2, Col4a3, Col4a4, Col14a1*), other ECM glycoproteins (*Ahsg, Atrnl1, Cilp, Ecm1, Fbln1/2, Fn1, Hmcn2, Mfap4/5, Thbs3/4, Tnxb*), metallopeptidases (*Adamts2, Adamts10, Adamtsl2*), regulators of ECM structure and organisation (*Loxl2, Pcolce2, Ssc5d*), small leucine-rich repeat proteoglycans (*Aspn, Bgn, Hspg2, Prelp*) and growth factors/signalling genes (*Ccn2, IgUbp7, Ltbp3/4, Ndnf*). Many of the proteins encoded by these genes (e.g. collagens, FN1, ADAMTS2, LTBP4) are secreted from fibroblasts, revealing a potential role for B55α in cardiomyocyte-fibroblast crosstalk. Cardiomyocytes serve an important paracrine function in regulating collagen production in fibroblasts.^75^ The mechanisms by which B55α in male cardiomyocytes influences fibroblast gene transcription and extracellular matrix composition requires further investigation.

We detected a similar increase in LV collagen content in female cKO hearts, despite no differences in expression of *Col1a1* or *Col3a1*, the predominant cardiac collagen isoforms. Expression of collagen IV isoforms (*Col4a3* and *Col4a4*) tended to be higher in female cKO vs FC hearts, but these changes alone cannot account for the increase in LV collagen by picrosirius red staining given that male cKO mice displayed greater increases in *Col4a3* and *Col4a4*, in addition to increased expression of several other collagen isoforms. It is possible that deletion of B55α in the female heart modulates the activity of enzymes involved in collagen degradation (e.g. TIMPs, MMPs, ADAMs), but investigation of this was beyond the scope of the current study. Of note, a previous study reported PP2A-dependent dephosphorylation of MMP-2 in isolated rat hearts subjected to ischemia/reperfusion, but the B subunit responsible for targeting PP2A catalytic activity to MMP-2 was not identified.^76^ Further work is required to understand B55α-dependent sex differences in the transcriptional and post-translational regulation of cardiac collagen composition.

Pathway analysis of downregulated genes in hearts of male cardiomyocyte-specific B55α knockout mice revealed an enrichment of pathways associated with the mitochondria and oxidative phosphorylation (see **Fig 6d** & **Supp Table 8**). A previous study reported differences in the expression of selected metabolic genes in hearts of mice with cardiomyocyte-specific deletion of the PP2A catalytic isoform PP2A-Cα between P1 and P11, but expression in the adult heart was not reported.^72^ As FoxO1 has previously been identified as a key regulator of mitochondrial enzymes in the heart,^45^ and B55α dephosphorylates FoxO1 in islet β-cells,^18^ we hypothesised that differences in the expression of mitochondrial and metabolic genes in B55α knockout hearts may be due to phosphorylation-mediated inactivation of FoxO1. However, we observed no differences in the phosphorylation of FoxO1 at the pertinent phosphosites between genotypes, suggesting a FoxO1-independent mechanism by which B55α regulates the expression of metabolic genes. Further work is required to identify the signalling mechanisms by which B55α regulates gene transcription in cardiomyocytes.

Finally, our findings may have implications for the development of PP2A-targeting small molecules, which are promising cancer therapeutics. PP2A is inactivated in numerous solid and haematological tumours, and compounds that increase PP2A activity (e.g. FTY720) reduce tumour burden in preclinical models.^77^ Conversely, PP2A inhibitors (e.g. LB100) have been shown to synergistically enhance the efficacy of other anticancer agents.^78^ A major challenge for developing more effective cancer therapies is to identify the specific PP2A complexes to be targeted.^79^ In this context, B55α and B56α target PP2A catalytic activity to different amino acid residues on the proto-oncogene c-Myc, with opposite effects on c-Myc stabilisation and oncogenic signalling.^43^ PP2A-B55α-mediated dephosphorylation of Thr58 leads to c-Myc stabilisation and tumorigenesis, while PP2A-B56α-mediated dephosphorylation of Ser62 leads to c-Myc destabilisation and tumour suppression, revealing distinct roles for different PP2A holoenzymes.^43^ PP2A B55 family members are emerging as important modulators of oncogenic signalling in numerous cancer settings.^80,81^ B55α has been identified as a tumour suppressor in breast and prostate cancers, leukemias, and lung and thyroid carcinomas, and promotes tumour growth in pancreatic cancer settings.^81^ Therapeutic approaches to activate or inhibit PP2A-B55α holoenzymes in specific cancer contexts need to consider the potential off-target effects of B55α modulation in the heart and other organs.

Our data show that partial loss of B55α in all cell types, or complete loss of B55α in cardiomyocytes, does not lead to pathological cardiac hypertrophy or dysfunction in mice in the absence of stress/injury. However, loss of B55α significantly altered the cardiac transcriptome in male mice and may predispose male hearts to pathology. Therefore, off-target cardiac effects of cancer therapeutics inhibiting B55α may be of concern in the male heart (and particularly in the aged male heart). Further research is required to investigate sex-specific responses to reduced B55α expression in settings of cardiac stress/injury, and if increasing B55α expression has any adverse effects on heart structure or function, or if it is cardioprotective.

In conclusion, this study provides the first characterisation of a PP2A B/B55 subfamily member in the mouse heart. Global disruption of the gene encoding B55α was embryonically lethal but did not adversely affect cardiac development. Loss of cardiomyocyte B55α did not impact overall heart size, atrial weight or LV dimensions in mice evaluated at 10-12 weeks of age. However, cardiomyocyte-specific deletion of B55α significantly altered the cardiac transcriptome in male mice, upregulating the expression of genes associated with the extracellular matrix and increasing LV collagen content, and downregulating the expression of genes associated with mitochondrial energy production. In contrast, few genes were differentially expressed in female hearts with and without cardiomyocyte B55α. In addition, mice with global heterozygous knockout of B55α had thinner LV walls and reduced mRNA, but not protein, expression of the lymphatic vessel marker *Lyve1*. These studies identify sex-specific functions for B55α in the heart under basal conditions and lay the foundation for future work investigating the role of B55α in settings of cardiac stress or injury.

## Acknowledgements

This work was supported by a National Heart Foundation Future Leader Fellowship to KLW (award ID 102539) and an Emerging Leader Fellowship from The Shine On Foundation. NMS was supported by an Australian Government Research Training Program Scholarship. JRM was supported by a National Health and Medical Research Council Senior Research Fellowship (grant ID 1078985) and Baker Fellowship (The Baker Foundation, Australia). This study utilised the Phenomics Australia Histopathology and Slide Scanning Service, University of Melbourne.

## Author contribution statement

NMS, HK, DGD, AJT & KLW performed experiments and data analysis. NMS, AJT & KLW prepared figures. VCG, KAS, AJAR, JRB, KMM & LMDD helped plan experiments and provided important intellectual input. DGD, LMDD, JRM & KLW contributed to student supervision. NMS & KLW wrote the manuscript. JRM & KLW developed the project and provided funding. All authors read and approved the final manuscript.

