## Supplementary material for "Sex-specific regulation of the cardiac transcriptome by the protein phosphatase 2A regulatory subunit B55α": Data Supplement

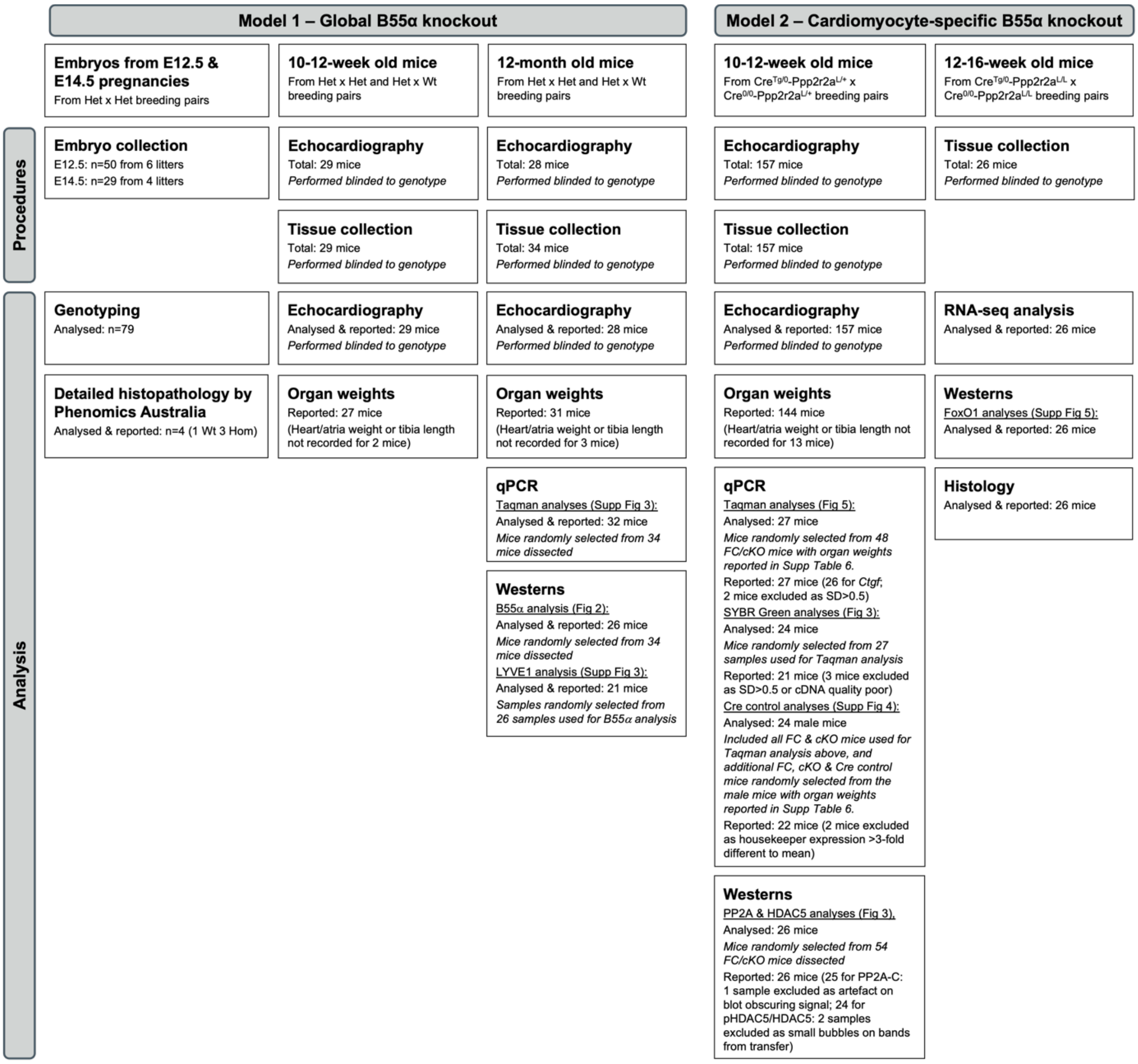


**Supp Fig 1: Animal numbers and data exclusions.**

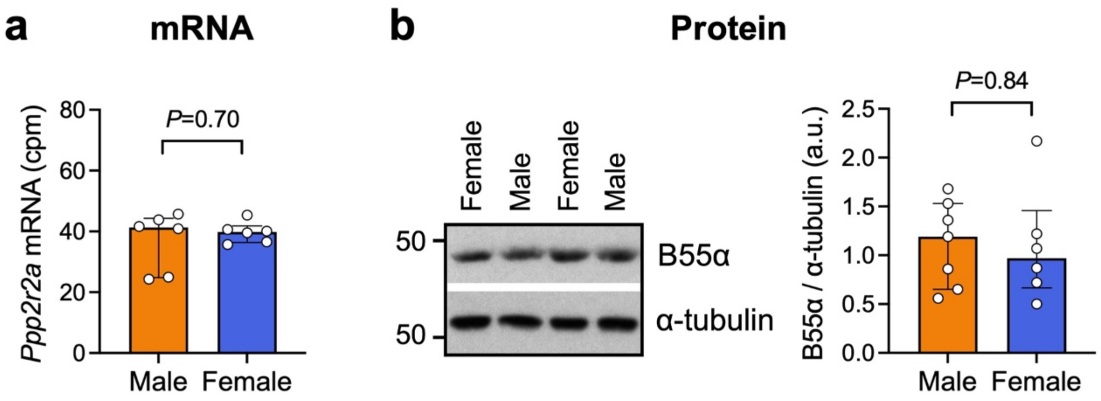


**Supp Fig 2: B55α mRNA and protein levels in male vs female control hearts.** a) B55α mRNA abundance (counts per million by RNA-seq) in ventricular tissue from male and female hearts (floxed controls from cardiomyocyte-specific B55α knockout model). Bars show median +/- IQR. Mann-Whitney U test. n=6/group. b) Representative Western blot and quantitation of B55α protein levels normalised to α-tubulin (arbitrary units, a.u.) in ventricular tissue from male and female hearts (wildtype controls from global B55α knockout model). Bars show median +/- IQR. Mann-Whitney U test. n=6-7 per group.

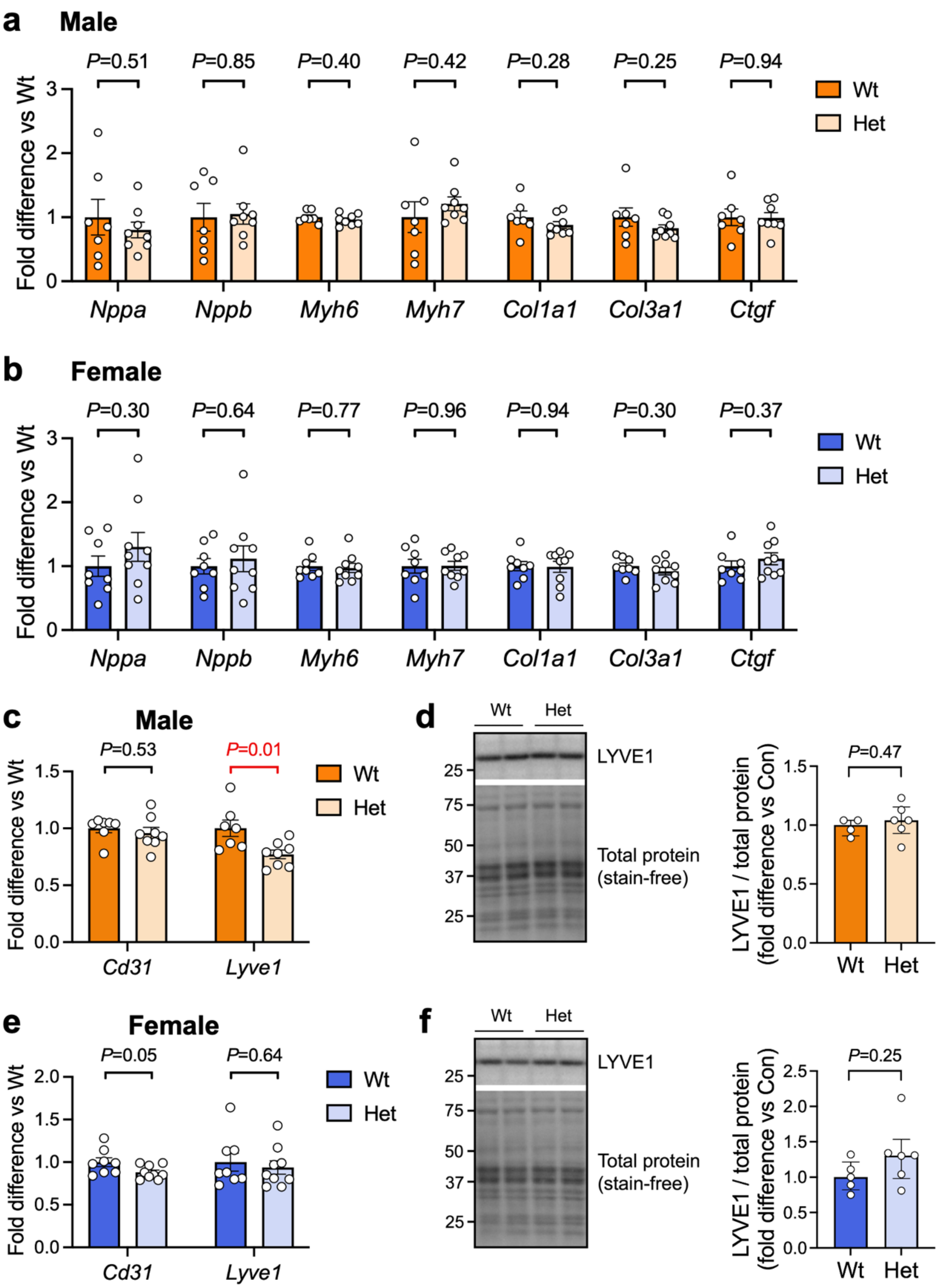


**Supp Fig 3: Cardiac gene and protein expression in mice with heterozygous expression of B55α at 12 months of age (global model).** RNA and protein were extracted from the ventricles of 12-month old male and female *Ppp2r2a* wildtype (Wt) and heterozygous (Het) mice. a, b) Quantitative PCR analysis of genes that are typically dysregulated in settings of pathological cardiac hypertrophy and fibrosis. *n*=7-9/group. Bars show mean +/- SEM. Unpaired t-tests. c, e) Quantitative PCR analysis of Cd31 and Lyve1 expression. n=7-9/group. Bars show mean +/- SEM. Unpaired t-tests. d, f) Representative Western blots and quantitation of LYVE1 normalised to total protein. n=4-6/group. Bars show median +/- IQR. Mann Whitney U tests.

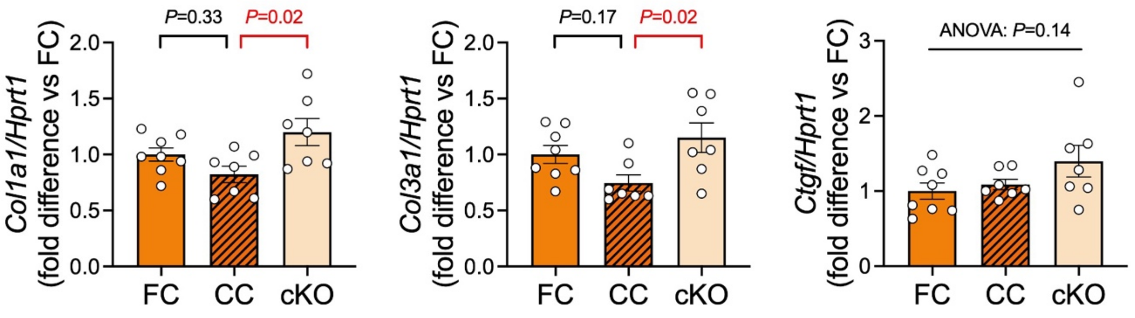


**Supp Fig 4: Comparison of collagen and *Ctgf* expression in floxed control (FC), Cre control (CC) and cardiomyocyte-specific B55α knockout (cKO) mice.** Quantitative PCR analysis of *Col1a1*, *Col3a1* and *Ctgf* expression in ventricular tissue from male mice. Data are expressed as fold difference relative to the FC group. Bars show mean +/- SEM. One-way ANOVA followed by Tukey's post-hoc tests. *n*=7-8 per group.

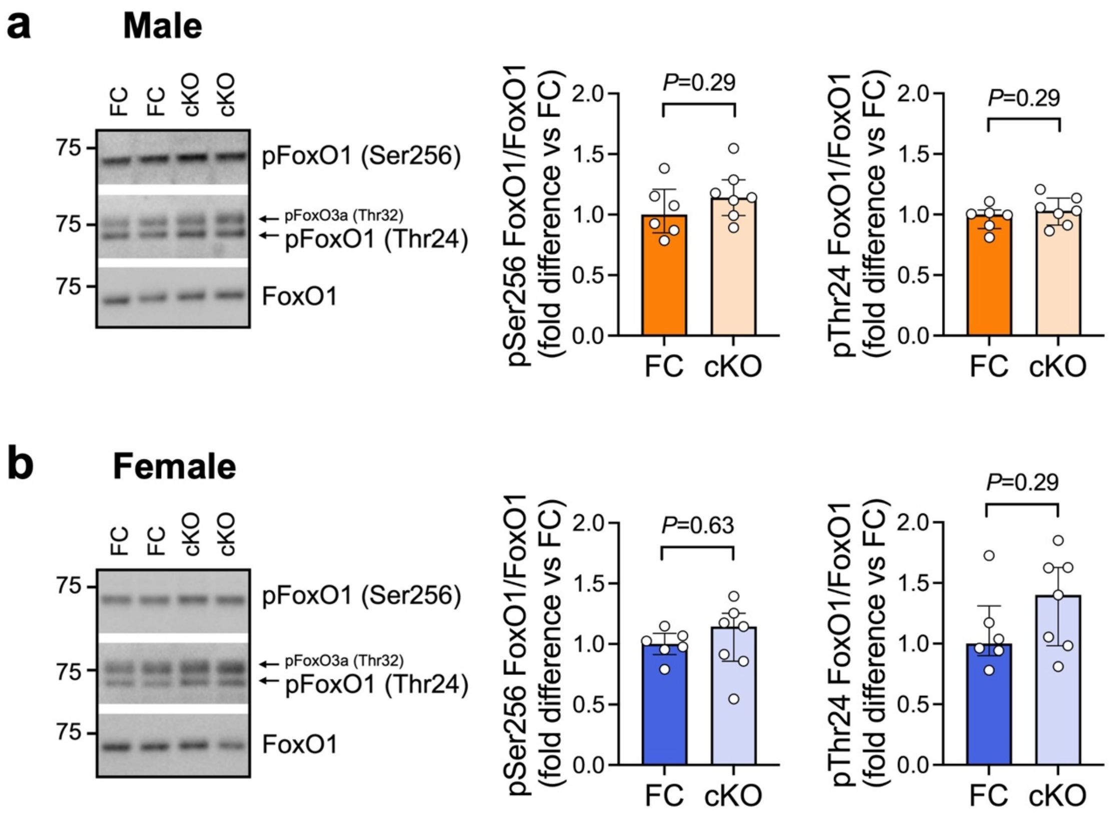


**Supp Fig 5: Phosphorylation of FoxO1 in cardiomyocyte-specific B55α knockout (cKO) and floxed control (FC) mice.** Representative Western blots and quantitation of FoxO1 phosphorylation at Ser256 and Thr24, normalised to total FoxO1, in a) male and b) female mice. Data are expressed as fold difference relative to the FC group. Bars show median +/- IQR. Mann-Whitney U tests. *n*=6-7 per group.

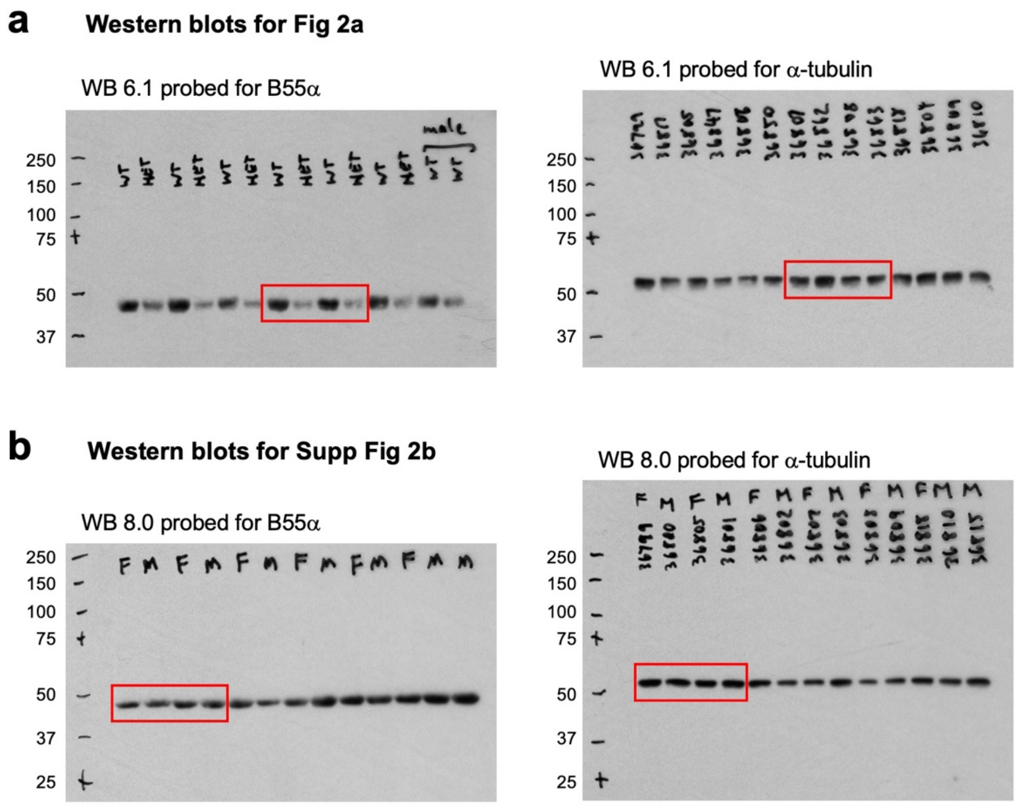


**Supp Fig 6: Uncropped Western blots, global B55α knockout mouse model.**
WT: wildtype, HET: heterozygote, F: female, M: male.

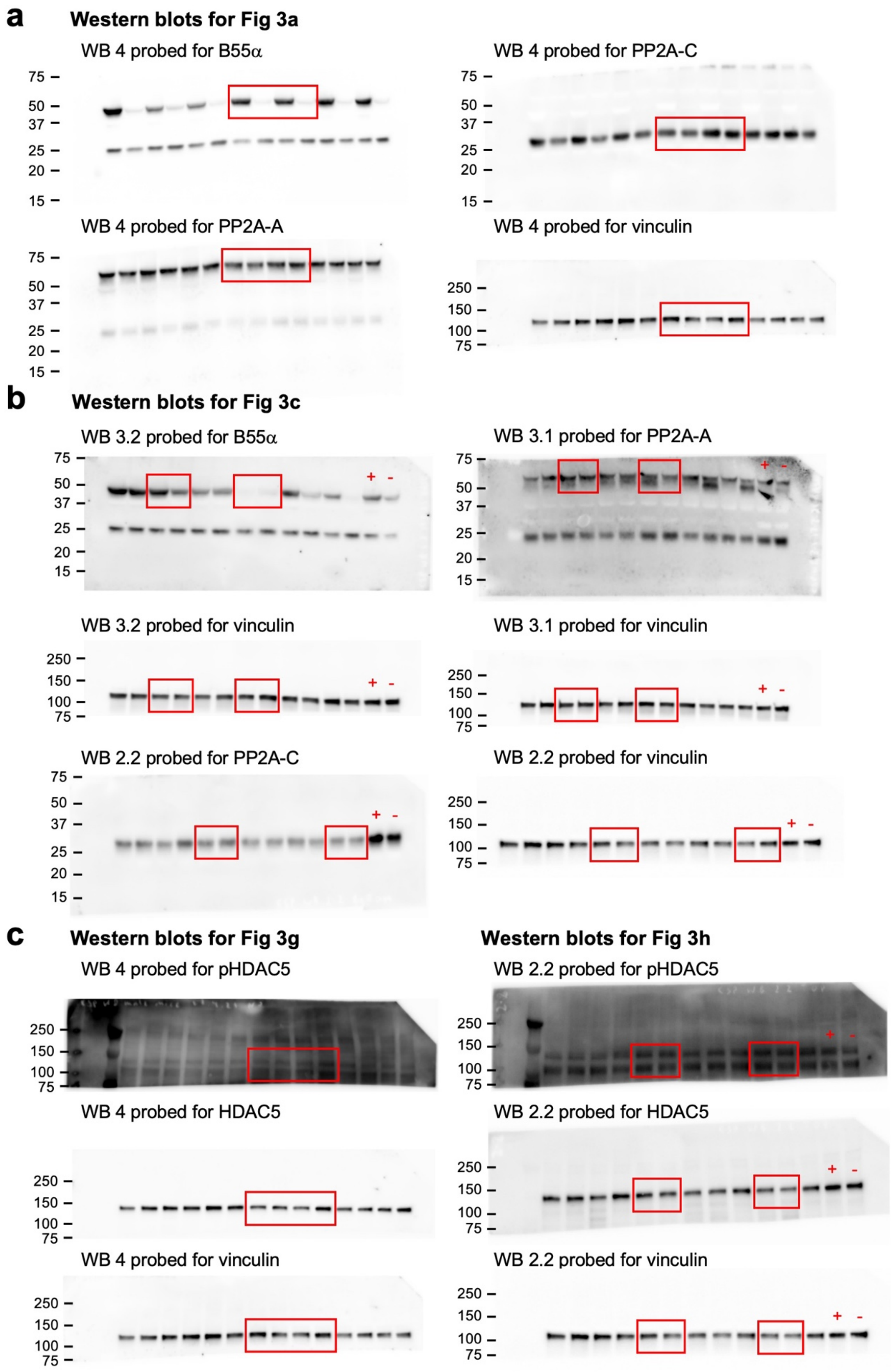


**Supp Fig 7: Uncropped Western blots, cardiomyocyte-specific B55α knockout mouse model.** WB: Western blot. LV lysates from a wildtype (+) and heterozygote (-) mouse from the global B55α knockout model were included on some blots to confirm specificity of the B55α antibody.

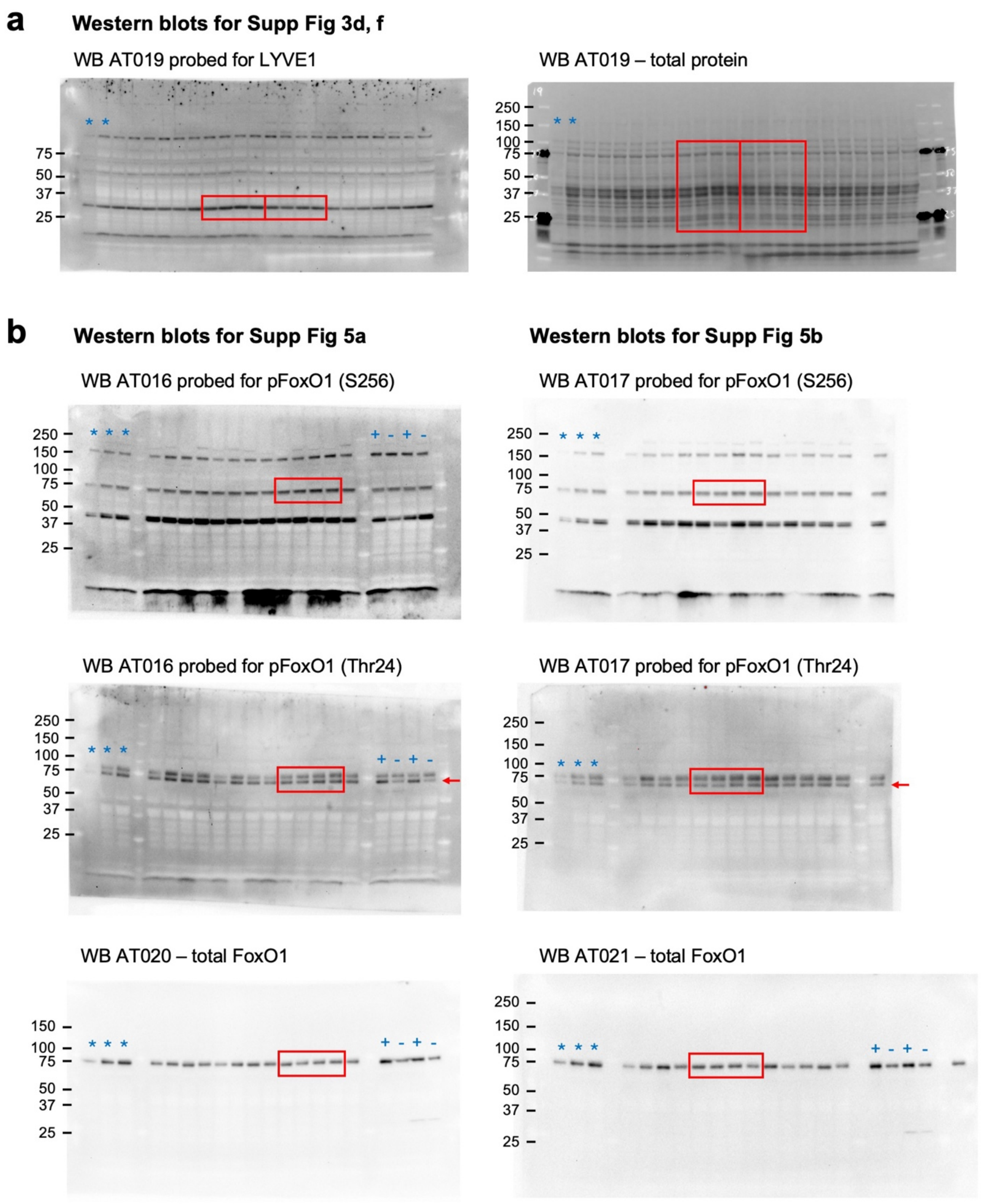


**Supp Fig 8: Uncropped Western blots, global and cardiomyocyte-specific B55α knockout mouse models.** LV lysates from control (+) and cardiomyocyte-specific FoxO1 knockout (-) mice (PMID 33577435) were included on some blots to confirm specificity of FoxO1 antibody signals. * denotes samples used to produce standard curves.

**Supp Table 1: Left ventricular morphology and function in 10-12 week old male and female mice with heterozygous expression of B55α (global knockout mouse model), assessed by M-mode echocardiography.** bpm: beats per minute; HR: heart rate; IVS: interventricular septum; LVPW: left ventricular posterior wall; LVID: left ventricular internal dimension at diastole (d) and systole (s); LV mass: estimated left ventricular mass; FS: fractional shortening. Data are median (Q1-Q3). Mann-Whitney U tests.

| ***10-12 weeks of age*** | **Wildtype (Wt)** | | **Heterozygous (Het)** | | ***P*-value** |
|---|---|---|---|---|---|
| ***MALES*** | **n=8** | | **n=8** | | |
| HR (bpm) | 490 | (458-500) | 472 | (466-483) | 0.34 |
| IVS (mm) | 0.62 | (0.48-0.68) | 0.59 | (0.52-0.67) | 0.98 |
| LVPW (mm) | 0.62 | (0.58-0.70) | 0.64 | (0.48-0.72) | 0.94 |
| LVID; d (mm) | 4.37 | (4.05-4.75) | 4.44 | (4.19-4.90) | 0.59 |
| LVID; s (mm) | 3.40 | (2.90-3.65) | 3.44 | (3.10-3.88) | 0.59 |
| FS (%) | 23 | (20-29) | 22 | (19-26) | 0.86 |
| LV mass (mg) | 98.6 | (88.1-105.6) | 91.9 | (83.9-115.3) | 0.88 |
| ***FEMALES*** | **n=7** | | **n=6** | | |
| HR (bpm) | 533 | (439-560) | 487 | (401-501) | 0.14 |
| IVS (mm) | 0.64 | (0.58-0.81) | 0.61 | (0.54-0.71) | 0.42 |
| LVPW (mm) | 0.64 | (0.58-0.73) | 0.59 | (0.45-0.71) | 0.35 |
| LVID; d (mm) | 3.92 | (3.84-4.44) | 4.47 | (3.97-4.58) | 0.31 |
| LVID; s (mm) | 2.93 | (2.71-3.72) | 3.25 | (2.87-3.33) | 0.84 |
| FS (%) | 25 | (23-29) | 27 | (24-30) | 0.27 |
| LV mass (mg) | 88.9 | (81.9-103.5) | 86.5 | (78.4-107.3) | 0.84 |

**Supp Table 2: Left ventricular morphology and function in 12-month old male and female mice with heterozygous expression of B55α (global knockout mouse model), assessed by M-mode echocardiography.** bpm: beats per minute; HR: heart rate; IVS: interventricular septum; LVPW: left ventricular posterior wall; LVID: left ventricular internal dimension at diastole (d) and systole (s); LV mass: estimated left ventricular mass; FS: fractional shortening. Data are median (Q1-Q3). Mann-Whitney U tests.

| ***12 months of age*** | **Wildtype (Wt)** | | **Heterozygous (Het)** | | ***P*-value** |
|---|---|---|---|---|---|
| ***MALES*** | **n=8** | | **n=8** | | |
| HR (bpm) | 500 | (484-528) | 483 | (463-535) | 0.46 |
| IVS (mm) | 1.02 | (0.95-1.08) | 0.94 | (0.88-0.98) | 0.04 |
| LVPW (mm) | 1.05 | (0.96-1.11) | 0.98 | (0.91-1.10) | 0.047 |
| LVID; d (mm) | 4.27 | (3.74-4.66) | 4.33 | (3.80-4.56) | 0.78 |
| LVID; s (mm) | 2.85 | (2.41-3.02) | 2.79 | (2.30-3.18) | 0.86 |
| FS (%) | 34 | (32-38) | 34 | (30-41) | 0.75 |
| LV mass (mg) | 183.8 | (153.4-228.4) | 170.6 | (132.0-176.0) | 0.20 |
| ***FEMALES*** | **n=6** | | **n=6** | | |
| HR (bpm) | 505 | (441-548) | 510 | (489-546) | 0.66 |
| IVS (mm) | 0.68 | (0.63-0.76) | 0.60 | (0.57-0.63) | 0.01 |
| LVPW (mm) | 0.64 | (0.60-0.66) | 0.63 | (0.60-0.67) | 0.91 |
| LVID (mm) | 4.39 | (4.21-4.53) | 4.50 | (4.34-4.67) | 0.29 |
| LVID, s (mm) | 3.21 | (2.86-3.35) | 3.40 | (2.98-3.48) | 0.29 |
| FS (%) | 26 | (25-35) | 26 | (24-32) | 0.91 |
| LV mass (mg) | 105.2 | (99.9-114.5) | 102.2 | (94.6-111.5) | 0.31 |

**Supp Table 3: Organ weights in 10-12-week old male and female mice with heterozygous expression of B55α (global knockout mouse model).** AW: atria weight, HW: heart weight, KidW: kidney weight, LivW: liver weight, LungW: lung weight, SpW: spleen weight, TL: tibia length. Data are median (Q1-Q3). Mann-Whitney U tests.

| ***10-12 weeks of age*** | **Wildtype (Wt)** | | **Heterozygous (Het)** | | ***P*-value** |
|---|---|---|---|---|---|
| ***MALES*** | **n=7** | | **n=7** | | |
| Tibia length (mm) | 16.1 | (16.0-16.3) | 15.7 | (15.6-16.0) | 0.004 |
| Body weight (g) | 26.5 | (26.4-28.3) | 27.5 | (24.9-28.7) | 0.69 |
| Heart (mg) | 126.4 | (120.1-130.5) | 122.8 | (111.5-137.3) | >0.99 |
| Atria (mg) | 7.0 | (6.6-7.3) | 6.2 | (5.7-6.6) | 0.03 |
| Lungs (mg) | 135.7 | (121.4-147.0) | 128.3 | (124.5-150.2) | 0.90 |
| Liver (mg) | 1251.0 | (1079.0-1383.0) | 1189.0 | (961.8-1328.0) | 0.63 |
| Kidney (mg) | 164.2 | (153.6-169.9) | 157.3 | (140.8-180.0) | 0.71 |
| Spleen (mg) | 74.7 | (69.9-79.4) | 68.2 | (63.1-86.9) | 0.49 |
| HW/TL (mg/mm) | 7.75 | (7.34-8.16) | 7.80 | (7.17-8.80) | 0.81 |
| AW/TL (mg/mm) | 0.43 | (0.41-0.46) | 0.40 | (0.37-0.42) | 0.09 |
| LungW/TL (mg/mm) | 8.38 | (7.59-8.96) | 8.15 | (8.03-9.39) | 0.62 |
| LivW/TL (mg/mm) | 77.23 | (67.25-84.70) | 76.12 | (61.91-84.05) | 0.63 |
| KidW/TL (mg/mm) | 10.26 | (9.57-10.62) | 9.86 | (9.08-11.54) | 0.81 |
| SpW/TL (mg/mm) | 4.68 | (4.34-4.89) | 4.36 | (4.04-5.54) | 0.70 |
| HW/BW (mg/g) | 4.77 | (4.36-4.88) | 4.51 | (4.33-4.98) | 0.90 |
| AW/BW (mg/g) | 0.27 | (0.24-0.28) | 0.23 | (0.22-0.25) | 0.07 |
| LungW/BW (mg/g) | 4.89 | (4.80-5.33) | 5.06 | (4.62-5.24) | 0.83 |
| LivW/BW (mg/g) | 44.79 | (42.83-50.33) | 42.77 | (39.35-46.83) | 0.53 |
| KW/BW (mg/g) | 6.19 | (5.87-6.44) | 5.89 | (5.59-6.20) | 0.46 |
| SW/BW (mg/g) | 2.78 | (2.67-2.91) | 2.61 | (2.41-3.04) | 0.49 |
| ***FEMALES*** | **n=7** | | **n=6** | | |
| Tibia length (mm) | 15.9 | (15.9-16.0) | 15.5 | (15.2-15.6) | 0.004 |
| Body weight (g) | 22.5 | (20.7-23.8) | 20.8 | (19.8-22.1) | 0.07 |
| Heart (mg) | 113.8 | (103.8-125.0) | 107.8 | (99.7-118.5) | 0.45 |
| Atria (mg) | 6.4 | (5.4-6.6) | 6.1 | (5.0-7.0) | 0.51 |
| Lungs (mg) | 128.9 | (126.2-138.6) | 129.8 | (123.1-133.5) | 0.84 |
| Liver (mg) | 935.6 | (878.2-1069.0) | 787.3 | (638.9-821.7) | 0.001 |
| Kidney (mg) | 131.6 | (126.7-139.0) | 118.6 | (112.6-136.8) | 0.10 |
| Spleen (mg) | 88.4 | (78.0-95.2) | 91.3 | (78.6-94.6) | 0.84 |
| HW/TL (mg/mm) | 7.08 | (6.55-7.86) | 7.03 | (6.44-7.62) | 0.84 |
| AW/TL (mg/mm) | 0.40 | (0.34-0.42) | 0.40 | (0.33-0.45) | 0.92 |
| LungW/TL (mg/mm) | 8.13 | (7.94-8.61) | 8.36 | (8.09-8.61) | 0.95 |
| LivW/TL (mg/mm) | 59.41 | (55.41-67.21) | 50.93 | (41.70-53.89) | 0.005 |
| KidW/TL (mg/mm) | 8.30 | (7.94-8.74) | 7.70 | (7.31-8.85) | 0.23 |
| SW/TL (mg/mm) | 5.55 | (4.92-5.99) | 5.91 | (5.12-6.10) | 0.53 |
| HW/BW (mg/g) | 5.00 | (4.63-5.41) | 5.46 | (4.57-5.73) | 0.63 |
| AW/BW (mg/g) | 0.29 | (0.25-0.29) | 0.31 | (0.23-0.34) | 0.44 |
| LungW/BW (mg/g) | 5.78 | (5.61-6.29) | 6.32 | (5.78-6.54) | 0.13 |
| LivW/BW (mg/g) | 42.84 | (38.48-47.11) | 36.70 | (32.45-38.50) | 0.01 |
| KW/BW (mg/g) | 6.02 | (5.32-6.28) | 5.99 | (5.16-6.63) | 0.95 |
| SW/BW (mg/g) | 3.90 | (3.72-4.27) | 4.29 | (3.87-4.56) | 0.30 |

**Supp Table 4: Organ weights in 12-month old male and female mice with heterozygous expression of B55α (global knockout mouse model).** AW: atria weight, BW: body weight, HW: heart weight, KidW: kidney weight, LivW: liver weight, LungW: lung weight, SpW: spleen weight, TL: tibia length. Data are median (Q1-Q3). Mann-Whitney U tests.

| ***12 months of age*** | **Wildtype (Wt)** | | **Heterozygous (Het)** | | ***P*-value** |
|---|---|---|---|---|---|
| ***MALES*** | **n=8** | | **n=8** | | |
| Tibia length (mm) | 17.2 | (17.1-17.5) | 17.1 | (16.8-17.2) | 0.08 |
| Body weight (g) | 44.3 | (37.8-45.4) | 44.3 | (38.7-47.0) | 0.80 |
| Heart (mg) | 195.8 | (178.2-222.5) | 183.9 | (160.0-195.7) | 0.13 |
| Atria (mg) | 10.3 | (7.9-14.2) | 8.2 | (7.1-9.6) | 0.07 |
| Lungs (mg) | 164.6 | (159.2-173.8) | 158.9 | (145.4-170.1) | 0.38 |
| Liver (mg) | 1528.0 | (1185.0-1761.0) | 1999.0 | (1558.0-2288.0) | 0.05 |
| Kidney (mg) | 228.8 | (200.6-257.9) | 197.5 | (184.0-210.7) | 0.08 |
| Spleen (mg) | 106.1 | (80.2-127.7) | 83.6 | (79.6-95.3) | 0.24 |
| HW/TL (mg/mm) | 11.17 | (10.32-12.95) | 10.80 | (9.54-11.63) | 0.38 |
| AW/TL (mg/mm) | 0.60 | (0.46-0.82) | 0.48 | (0.42-0.56) | 0.11 |
| LungW/TL (mg/mm) | 9.54 | (9.29-10.10) | 9.41 | (8.49-9.93) | 0.57 |
| LivW/TL (mg/mm) | 89.05 | (69.71-100.80) | 117.60 | (92.41-131.60) | 0.05 |
| KW/TL (mg/mm) | 13.10 | (11.52-15.00) | 11.58 | (10.86-12.32) | 0.13 |
| SW/TL (mg/mm) | 6.19 | (4.59-7.42) | 4.91 | (4.73-5.49) | 0.24 |
| HW/BW (mg/g) | 4.45 | (4.25-5.03) | 4.25 | (4.02-4.84) | 0.34 |
| AW/BW (mg/g) | 0.24 | (0.20-0.33) | 0.18 | (0.17-0.23) | 0.14 |
| LungW/BW (mg/g) | 3.90 | (3.64-4.33) | 3.84 | (3.42-4.59) | 0.80 |
| LivW/BW (mg/g) | 34.52 | (32.17-39.06) | 46.12 | (43.18-49.22) | <0.001 |
| KW/BW (mg/g) | 4.98 | (4.59-6.51) | 4.71 | (4.31-5.35) | 0.24 |
| SW/BW (mg/g) | 2.56 | (1.78-2.87) | 2.02 | (1.95-2.39) | 0.44 |
| ***FEMALES*** | **n=8** | | **n=7** | | |
| Tibia length (mm) | 16.9 | (16.7-17.0) | 16.7 | (16.6-17.0) | 0.20 |
| Body weight (g) | 33.3 | (30.4-35.4) | 36.8 | (31.5-40.3) | 0.40 |
| Heart (mg) | 126.1 | (118.3-133.7) | 126.2 | (116.2-133.3) | 0.78 |
| Atria (mg) | 6.1 | (5.1-6.7) | 5.9 | (5.3-6.70) | 0.67 |
| Lungs (mg) | 152.8 | (145.2-160.0) | 143.3 | (139.9-147.7) | 0.05 |
| Liver (mg) | 1180.0 | (1025.0-1454.0) | 1357.0 | (1174.0-1600.0) | 0.46 |
| Kidney (mg) | 154.0 | (139.2-171.3) | 143.3 | (129.3-159.4) | 0.14 |
| Spleen (mg) | 85.0 | (71.6-95.6) | 80.9 | (71.5-98.7) | >0.99 |
| HW/TL (mg/mm) | 7.50 | (7.05-7.96) | 7.54 | (6.96-7.85) | 0.61 |
| AW/TL (mg/mm) | 0.36 | (0.30-0.40) | 0.35 | (0.32-0.39) | 0.71 |
| LungW/TL (mg/mm) | 9.15 | (8.62-9.39) | 8.49 | (8.40-8.69) | 0.07 |
| LivW/TL (mg/mm) | 70.18 | (60.29-86.57) | 80.53 | (71.27-95.97) | 0.46 |
| KidW/TL (mg/mm) | 9.20 | (8.25-10.06) | 8.50 | (7.79-9.38) | 0.15 |
| SW/TL (mg/mm) | 5.03 | (4.27-5.65) | 4.76 | (4.31-5.92) | 0.96 |
| HW/BW (mg/g) | 3.71 | (3.54-4.23) | 3.46 | (3.13-3.65) | 0.12 |
| AW/BW (mg/g) | 0.17 | (0.15-0.22) | 0.16 | (0.15-0.18) | 0.48 |
| LungW/BW (mg/g) | 4.44 | (4.11-5.09) | 4.14 | (3.56-4.37) | 0.18 |
| LivW/BW (mg/g) | 39.09 | (31.58-41.10) | 39.75 | (36.84-41.78) | 0.54 |
| KW/BW (mg/g) | 4.63 | (4.41-4.87) | 4.01 | (3.84-4.22) | <0.001 |
| SW/BW (mg/g) | 2.46 | (2.33-2.89) | 2.27 | (2.16-2.51) | 0.40 |

**Supp Table 5: Left ventricular morphology and function in 10-12-week old male and female mice with cardiomyocyte-specific knockout of B55α, assessed by M-mode echocardiography.** bpm: beats per minute; HR: heart rate; IVS: interventricular septum; LVPW: left ventricular posterior wall; LVID: left ventricular internal dimension at diastole (d) and systole (s); LV mass: estimated left ventricular mass; FS: fractional shortening. Data are median (Q1-Q3). Kruskal-Wallis tests.

| ***Males*** | **Cre$^{0/0}$-Ppp2r2a$^{+/+}$ (wildtype control)** | | **Cre$^{0/0}$-Ppp2r2a$^{L/+}$ (half floxed control)** | | **Cre$^{0/0}$-Ppp2r2a$^{L/L}$ (floxed control, FC)** | | **Cre$^{Tg/0}$-Ppp2r2a$^{+/+}$ (Cre control)** | | **Cre$^{Tg/0}$-Ppp2r2a$^{L/+}$ (heterozygote)** | | **Cre$^{Tg/0}$-Ppp2r2a$^{L/L}$ (knockout, cKO)** | | ***P*-value** |
|---|---|---|---|---|---|---|---|---|---|---|---|---|---|
| n | 13 | | 13 | | 13 | | 13 | | 13 | | 13 | | |
| HR (bpm) | 529 | (507-556) | 548 | (500-564) | 524 | (502-550) | 536 | (446-553) | 528 | (487-552) | 541 | (509-582) | 0.73 |
| IVS (mm) | 0.84 | (0.80-0.88) | 0.87 | (0.82-0.91) | 0.83 | (0.77-0.87) | 0.82 | (0.78-0.88) | 0.80 | (0.74-0.85) | 0.80 | (0.73-0.83) | 0.23 |
| LVID; d (mm) | 4.06 | (3.68-4.22) | 3.86 | (3.45-4.27) | 3.93 | (3.82-4.27) | 4.01 | (3.34-4.35) | 4.14 | (3.75-4.18) | 4.19 | (3.95-4.60) | 0.30 |
| LVPW (mm) | 0.84 | (0.74-0.92) | 0.87 | (0.81-0.99) | 0.81 | (0.76-0.87) | 0.81 | (0.80-0.93) | 0.78 | (0.75-0.88) | 0.76 | (0.74-0.82) | 0.08 |
| LVID; s (mm) | 2.57 | (2.20-2.95) | 2.62 | (2.23-3.19) | 2.78 | (2.56-2.86) | 2.86 | (2.06-3.13) | 2.88 | (2.36-3.02) | 2.91 | (2.68-3.36) | 0.30 |
| FS (%) | 37 | (31-42) | 32 | (26-36) | 33 | (28-36) | 32 | (25-36) | 32 | (28-38) | 30 | (26-31) | 0.11 |
| LV mass (mg) | 123.9 | (114.3-133.0) | 128.5 | (107.1-132.5) | 118.6 | (107.3-132.2) | 115.0 | (106.4-136.2) | 117.7 | (104.9-132.9) | 122.2 | (114.8-135.0) | 0.88 |
| ***Females*** | **Cre$^{0/0}$-Ppp2r2a$^{+/+}$ (wildtype control)** | | **Cre$^{0/0}$-Ppp2r2a$^{L/+}$ (half floxed control)** | | **Cre$^{0/0}$-Ppp2r2a$^{L/L}$ (floxed control, FC)** | | **Cre$^{Tg/0}$-Ppp2r2a$^{+/+}$ (Cre control)** | | **Cre$^{Tg/0}$-Ppp2r2a$^{L/+}$ (heterozygote)** | | **Cre$^{Tg/0}$-Ppp2r2a$^{L/L}$ (knockout, cKO)** | | ***P*-value** |
| n | 15 | | 13 | | 13 | | 12 | | 13 | | 13 | | |
| HR (bpm) | 486 | (453-509) | 522 | (497-537) | 497 | (443-514) | 482 | (432-512) | 504 | (446-529) | 529 | (470-549) | 0.16 |
| IVS (mm) | 0.77 | (0.71-0.83) | 0.77 | (0.70-0.84) | 0.80 | (0.71-0.86) | 0.75 | (0.66-0.77) | 0.80 | (0.74-0.88) | 0.81 | (0.72-0.85) | 0.59 |
| LVID; d (mm) | 4.03 | (3.49-4.17) | 3.98 | (3.69-4.11) | 3.96 | (3.66-4.14) | 3.74 | (3.47-4.02) | 3.72 | (3.51-4.00) | 3.75 | (3.59-4.12) | 0.36 |
| LVPW (mm) | 0.77 | (0.70-0.85) | 0.77 | (0.76-0.81) | 0.77 | (0.64-0.83) | 0.71 | (0.66-0.82) | 0.77 | (0.74-0.86) | 0.75 | (0.68-0.83) | 0.56 |
| LVID; s (mm) | 2.77 | (2.36-3.03) | 2.66 | (2.48-2.95) | 2.71 | (2.45-3.02) | 2.50 | (2.26-2.84) | 2.34 | (2.16-2.69) | 2.62 | (2.46-2.85) | 0.16 |
| FS (%) | 31 | (30-34) | 29 | (28-37) | 29 | (26-36) | 34 | (31-37) | 35 | (32-40) | 30 | (30-31) | 0.06 |
| LV mass (mg) | 106.6 | (101.0-116.5) | 105.3 | (99.80-119.9) | 98.7 | (94.15-126.1) | 93.2 | (79.73-105.7) | 107.7 | (96.20-112.5) | 106.4 | (90.2-117.8) | 0.15 |

**Supp Table 6: Organ weights in 10-12-week old male and female mice with cardiomyocyte-specific knockout of B55α.**
AW: atria weight, BW: body weight, HW: heart weight, LW: lung weight, TL: tibia length. Data are median (Q1-Q3). Kruskal-Wallis tests.

| ***Males*** | **$Cre^{0/0}$-$Ppp2r2a^{+/+}$ (wildtype control)** | **$Cre^{0/0}$-$Ppp2r2a^{L/+}$ (half floxed control)** | **$Cre^{0/0}$-$Ppp2r2a^{L/L}$ (floxed control, FC)** | **$Cre^{Tg/0}$-$Ppp2r2a^{+/+}$ (Cre control)** | **$Cre^{Tg/0}$-$Ppp2r2a^{L/+}$ (heterozygote)** | **$Cre^{Tg/0}$-$Ppp2r2a^{L/L}$ (knockout, cKO)** | ***P*-value** |
|---|---|---|---|---|---|---|---|
| n | 12 | 12 | 12 | 11 (10 for lung weight) | 12 | 12 | |
| TL (mm) | 16.7 (16.4-16.8) | 16.6 (16.3-16.7) | 16.8 (16.3-17.2) | 16.5 (16.1-17.0) | 16.6 (16.4-17.0) | 16.7 (16.4-17.1) | 0.53 |
| BW (g) | 29.2 (27.4-31.4) | 28.6 (26.8-29.7) | 29.9 (28.3-31.8) | 27.4 (26.4-30.3) | 29.3 (26.8-31.3) | 30.7 (28.4-34.4) | 0.24 |
| Heart (mg) | 145.1 (137.6-157.2) | 138.4 (125.6-149.2) | 144.3 (140.5-155.3) | 138.2 (126.3-142.5) | 144.2 (121.2-160.6) | 148.8 (135.6-155.8) | 0.49 |
| Atria (mg) | 7.8 (6.3-8.6) | 7.9 (6.5-8.7) | 7.4 (6.8-8.8) | 7.2 (6.0-8.5) | 7.7 (6.9-8.5) | 7.6 (6.5-8.1) | 0.99 |
| Lungs (mg) | 137.8 (131.5-149.7) | 142.0 (126.5-150.1) | 144.6 (138.9-149.2) | 126.9 (118.7-135.8) | 134.1 (127.7-152.1) | 140.7 (131.1-150.2) | 0.18 |
| HW/TL (mg/mm) | 8.81 (8.17-9.33) | 8.27 (7.60-8.91) | 8.70 (8.36-9.06) | 8.15 (7.68-8.79) | 8.39 (7.40-9.72) | 8.80 (8.24-9.21) | 0.61 |
| AW/TL (mg/mm) | 0.46 (0.38-0.53) | 0.47 (0.40-0.53) | 0.44 (0.40-0.53) | 0.45 (0.38-0.52) | 0.46 (0.43-0.50) | 0.45 (0.39-0.49) | 0.99 |
| LW/TL (mg/mm) | 8.32 (7.90-9.01) | 8.58 (7.55-9.02) | 8.55 (8.24-8.82) | 7.77 (7.05-8.28) | 8.10 (7.64-9.14) | 8.28 (7.90-9.10) | 0.31 |
| ***Females*** | **$Cre^{0/0}$-$Ppp2r2a^{+/+}$ (wildtype control)** | **$Cre^{0/0}$-$Ppp2r2a^{L/+}$ (half floxed control)** | **$Cre^{0/0}$-$Ppp2r2a^{L/L}$ (floxed control, FC)** | **$Cre^{Tg/0}$-$Ppp2r2a^{+/+}$ (Cre control)** | **$Cre^{Tg/0}$-$Ppp2r2a^{L/+}$ (heterozygote)** | **$Cre^{Tg/0}$-$Ppp2r2a^{L/L}$ (knockout, cKO)** | ***P*-value** |
| n | 13 | 13 | 12 | 11 | 12 | 12 | |
| TL (mm) | 16.2 (16.1-16.5) | 16.4 (16.1-16.7) | 16.3 (16.0-16.8) | 16.5 (16.2-16.7) | 16.3 (15.9-16.5) | 16.2 (16.0-16.5) | 0.72 |
| BW (g) | 24.4 (23.1-25.9) | 23.9 (21.8-25.4) | 23.5 (22.3-26.4) | 22.8 (21.3-23.1) | 22.7 (20.6-25.2) | 22.0 (20.8-25.3) | 0.19 |
| Heart (mg) | 110.5 (106.8-124.3) | 113.2 (101.7-120.2) | 109.5 (99.9-113.7) | 106.1 (101.4-115.6) | 107.0 (96.3-118.7) | 105.9 (104.5-110.8) | 0.55 |
| Atria (mg) | 5.5 (4.7-7.4) | 5.9 (5.0-6.7) | 5.7 (4.7-6.6) | 5.0 (4.8-5.6) | 4.7 (3.9-5.7) | 5.5 (4.8-6.5) | 0.27 |
| Lungs (mg) | 132.5 (114.5-136.7) | 129.1 (123.1-136.2) | 124.3 (118.2-129.0) | 121.0 (110.9-130.1) | 125.1 (110.9-132.4) | 121.8 (116.0-130.0) | 0.39 |
| HW/TL (mg/mm) | 6.86 (6.63-7.63) | 6.79 (6.30-7.33) | 6.60 (6.15-6.94) | 6.54 (6.11-6.92) | 6.54 (6.01-7.18) | 6.58 (6.42-6.78) | 0.37 |
| AW/TL (mg/mm) | 0.35 (0.29-0.45) | 0.37 (0.30-0.41) | 0.35 (0.30-0.40) | 0.31 (0.28-0.35) | 0.29 (0.25-0.36) | 0.36 (0.31-0.40) | 0.27 |
| LW/TL (mg/mm) | 8.15 (7.22-8.27) | 7.90 (7.40-8.23) | 7.59 (7.00-7.84) | 7.24 (6.61-8.02) | 7.72 (6.97-8.14) | 7.56 (7.23-7.90) | 0.32 |

**Supp Table 7: Differential gene expression analysis in male and female cardiomyocyte-specific B55α knockout mice vs control.** Tissue was collected at 12-16 weeks of age. *n*=6-7/group. False discovery rate (FDR) was calculated using Benjamini-Hochberg method and genes with an FDR<0.05 were considered differentially expressed.

*See Excel file titled: “Supp Table 7 – DEG cKO vs Con.xlsx”*

**Supp Table 8: Gene Ontology enrichment analyses (Biological Process, Cellular Component & Molecular Function) on differentially expressed genes in male cardiomyocyte-specific B55α knockout mice vs control.** Tissue was collected at 12-16 weeks of age. *n*=6-7/group.

*See Excel file titled: “Supp Table 8 – GO pathway enrichment, male cKO vs Con.xlsx”.*
